# Coordinated variation in the immune systems and microbiomes of healthy humans is linked to tonic interferon states

**DOI:** 10.1101/2024.11.27.625750

**Authors:** Joel Babdor, Ravi K. Patel, Brittany Davidson, Cecilia Noecker, Maha Rahim, Jordan E. Bisanz, Iliana Tenvooren, Diana Marquez, Maria Calvo, Vrinda Johri, Elizabeth McCarthy, Avneet Shaheed, Christina Ekstrand, Allison M. Weakley, Feiqiao B. Yu, Oliver Fiehn, Peter J. Turnbaugh, Alexis J. Combes, Gabriela K. Fragiadakis, Matthew H. Spitzer

## Abstract

Human immune systems are highly variable, with most variation attributable to non-genetic sources. The gut microbiome crucially shapes the immune system; however, its relationship with the baseline immune states of healthy humans remains incompletely understood. Therefore, we performed multi-omic profiling of 110 healthy participants through the ImmunoMicrobiome study. A factor-based integrative approach identified coordinated variation, revealing that the tonic interferon response was amongst the most variable immune features in healthy participants. Microbiome composition, pathways, and stool metabolites varied concomitantly with interferon response pathways. Distinct transcriptional programs involving inflammation and TGF-β in SIGLEC-1^high^ monocytes and CD69^high^ activated MAIT and NK cells were representative of these programs. Our study provides extensive data to examine the relationship between the immune states and microbiomes of healthy individuals at steady state, which paves the way for delineating inter-individual differences relevant for disease susceptibility and responses to therapy.

## Introduction

Inter-individual immune variation has broad implications for human health and disease. Population immunology studies have begun identifying drivers of immune variability, revealing that environmental factors likely outweigh genetic contributions.^1–8^ Understanding these drivers is challenging because of their complexity and intricacy but could provide insights into disease susceptibility, severity, and responsiveness to immunomodulatory therapies.^9^ Notably, immune-modulating interventions exhibit variable responses across patients in vaccination,^10,11^ cancer immunotherapy,^12^ and immunosuppressive treatments for autoimmune diseases.^13^

The gut microbiome has emerged as an important factor influencing immune responses. In cancer, human studies found microbiome associations with clinical outcomes of patients treated with checkpoint inhibitors.^14^ In autoimmunity, the microbiome and metabolites produced by commensal microbes also impact treatment response.^15,16^ In vaccination, the microbiota can act as an adjuvant, boosting immunogenicity of interventions.^17–20^ These studies suggest that the microbiome and molecules they produce could shape patients’ baseline immune states and influence immune responses to treatment. Animal models have revealed causal relationships between the microbiome and immune cell development and activity.^21,22^ Nevertheless, the relationship between microbiome features and variations in immune states among healthy humans at steady state remains incompletely understood.

One example is the regulation of baseline or tonic interferon (IFN) signaling, evidenced by the reduced expression of IFN-stimulated genes (ISGs) and impaired antiviral immune responses in microbiome-deficient mice.^19,23,24^ Indeed, IFNs are produced at low levels in homeostasis, and sensing of IFNs results in tonic signaling critical for antimicrobial immune responses^25–27^ as well as shaping the immune landscape^28^ and immune activity.^23^ Whether tonic-IFN signaling varies across healthy humans, and whether it is linked to the human gut microbiome, remains incompletely understood.

In this study we set out to investigate immunological variation across healthy individuals and to interrogate whether and which microbiome features are associated with these immune states. To do so, we generated multimodal profiling data of the immune system, the microbiome and the metabolome as a resource for the field. We used a factor-based integrative approach to capture co-varying features across the cohort. Combining our dataset with this method, we identified two major axes of immune variability in healthy individuals involving IFN response programs. While one axis was associated with gut microbiome features, the other was independent. The microbiome-associated axis of immune variation specifically captured tonic-IFN response gene expression along with differences in abundance and phenotypes of SIGLEC-1^high^ monocytes, activated CD8 and CD4 T cell subsets, and activated CD69^high^ NK and MAIT cell subsets. Validation datasets indicated that the microbiome-associated tonic-IFN signature was associated with increased vaccine responses but reduced cancer immunotherapy responses, suggesting potential applications of these immune states and biomarkers in personalized medicine.

## Results

### Immune and microbiome variation in a cohort of healthy individuals

To study variation in the immune system and the microbiome in humans at steady state, we recruited 110 healthy participants within the greater San Francisco Bay Area to establish the ImmunoMicrobiome cohort (**Fig. 1A**). Participants were all healthy adults of predominantly non-Hispanic White and Asian ancestries (**Fig. 1B, Methods**). We collected blood and stool specimens as well as survey information related to general health, diet, and medical history. We performed immune profiling of peripheral blood mononuclear cells (PBMCs), quantified circulating factors in plasma, and generated microbiome profiling data on DNA extracted from stool (**Fig. 1C**). We additionally generated mass spectrometry metabolomics data from stool and plasma. We preprocessed each dataset (Methods) to extract molecular and cellular features (**Table S1**), the distribution of which was assessed across the cohort.

**Figure 1.**
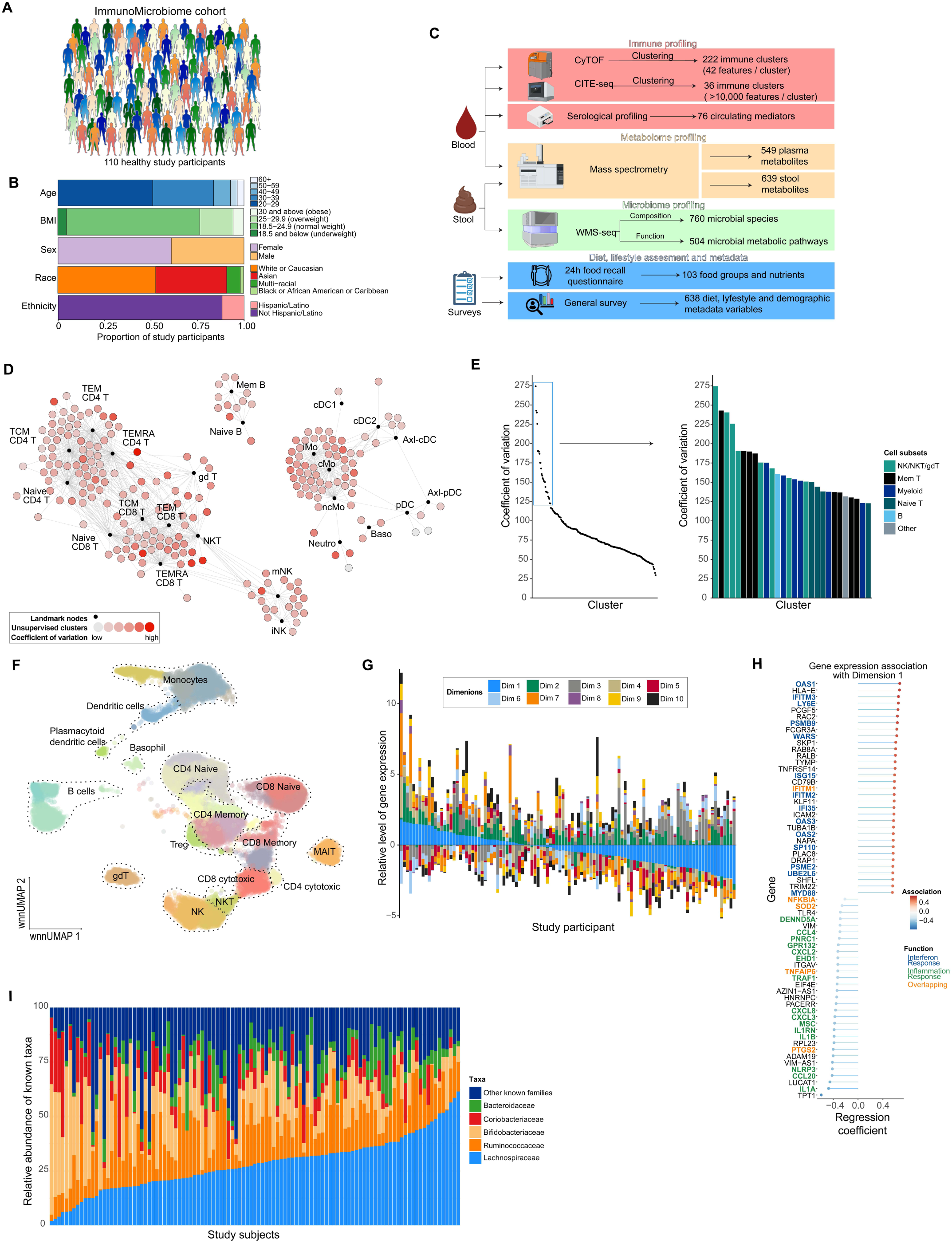
Immune and microbiome variation in a cohort of healthy individuals. A. ImmunoMicrobiome cohort overview. B. ImmunoMicrobiome cohort demographic breakdown. C. Schematic of the data generation strategy. D. Scaffold map of CyTOF unsupervised immune cell clusters across the cohort. Clusters are colored by their coefficient of variation calculated on cluster frequency across the cohort. Landmark nodes (black dots) represent defined immune populations used as a reference for the mapping. E. Dot plot of the coefficient of variation of CyTOF clusters across the cohort. The most variable clusters are described in a barchart. F. UMAP visualization of CITE-Seq data. Annotation and dotted line shapes indicate broad cell subsets. Single-cells are colored by clusters to indicate the diversity of subsets captured by the clustering approach within each cell subsets. Cluster level annotation is detailed in Fig. S1A-B. G. Transcriptional variation in monocytes across the cohort. The first ten dimensions obtained by principal component analysis and harmony batch correction on CITE-Seq gene expression data are shown. Study participant-specific dimensions’ coordinates are the average (means) of the coordinates of their single-cells. Participants are ordered by their coordinates for Dimension 1. H. Top genes significantly associated with Dimension 1 (adj.p < 0.1) in monocytes. Linear model is used to identify genes whose expressions are associated with Dimension 1. Genes are colored by presence in hallmark gene signatures. Green, inflammation (M5932 and M5890); blue, interferon response (M5911 and M5913); orange, overlap. I. Relative abundance of the top 5 most represented microbiome families across study participants.

We used single-cell mass cytometry (CyTOF) to evaluate immune cell population frequencies and their variation across the cohort. We annotated unsupervised clusters of cells based on expression of key markers (**Supplemental Methods**), and visualized their variation in relative abundance on a forced-directed graph (**Fig. 1D**).^29^ While most clusters were variable across individuals, certain NKT, NK, and T cell clusters showed the highest variation (**Fig. 1D-E**).

We used cellular indexing of transcriptomes and epitopes by sequencing (CITE-seq) to evaluate variation in gene and protein expression of immune cell subsets across the cohort. Single cells were clustered and annotated based on both extracellular protein expression and gene expression, identifying cell clusters that comprised established immune cell types (**Fig. 1F, S1A-B**). The gene expression matrix of each cell type was subjected to independent dimensionality reduction to assess the major transcriptional axes of variation across the cohort. As an example, in monocytes, we visualized variance in gene expression across the cohort in the top 10 dimensions (**Fig. 1G**). Dimension 1 was positively associated with genes involved in interferon response but negatively associated with genes involved in inflammation response (**Fig. 1H**). Variation in gene expression of a similar scale was also found for the other cell types across the cohort.

In parallel, we investigated variation in microbiome composition using metagenomic sequencing. We identified microbial clades at various taxonomic levels. As expected, the majority of species identified were part of 4 major phyla (**Fig. S1C**) and showed differences in diversity across the cohort (**Fig. S1D**). While the top 5 most abundant families covered more than 50% of the known taxa in most study participants, the relative abundance of each family varied widely across individuals (**Fig. 1I**). This variation of composition was also visible at the species level (**Fig. S1E**). These observations were broadly consistent with the major taxa and abundance variation found in the Human Microbiome Project,^30^ which sampled a larger population of healthy individuals.

Together, these high-level analyses highlight substantial inter-individual variation across immune and microbiome composition, as well as gene expression of key immune effectors, in a cohort of healthy individuals.

### Integrative multi-omic analysis identifies microbiome-associated immune variation in interferon and inflammation responses

To identify biological features that vary concomitantly between immune and microbiome systems, we used a multi-omic approach to integrate blood profiling, stool profiling and dietary recall data. We applied multi-omic factor analysis (MOFA)^31^ to generate factors composed of coordinated features across modalities that capture distinct biological variation across the cohort. This approach produced factor scores for each study participant for each factor, along with feature weights indicating the strength of association of features with each factor (**Fig. S2A**). Factors were numbered by variance explained. We identified features significantly associated with each factor using linear models (Methods) and evaluated the number and proportion of associated features from each modality with each factor (**Fig. 2A**). While factor 1 captured significant variation in immune cell gene expression, it was not significantly associated with stool microbiome or metabolome features (**Fig. 2A-B**). In contrast, factor 3 captured substantial variation in the immune system, microbiome species composition, microbiome pathway abundances, and metabolites. Among the top factors, factor 3 also captured the most variation in the relative abundance of immune cell subsets in both CyTOF and CITE-Seq data. Principal component analysis (PCA) of species-level microbiome composition data identified a significant correlation between MOFA factor 3 with PC2, as did PCA of CITE-Seq gene expression (**Fig. 2C, S2B**). In comparison, factor 1 was associated with CITE-Seq gene expression PCs but not with microbial composition PCs (**Fig. 2D, Fig. S2C**), validating our observations of these factors. We therefore refer to factor 1 as the independent immune variation (IIV) and to factor 3 as the immune and microbiome concomitant variation (IMCV).

**Figure 2.**
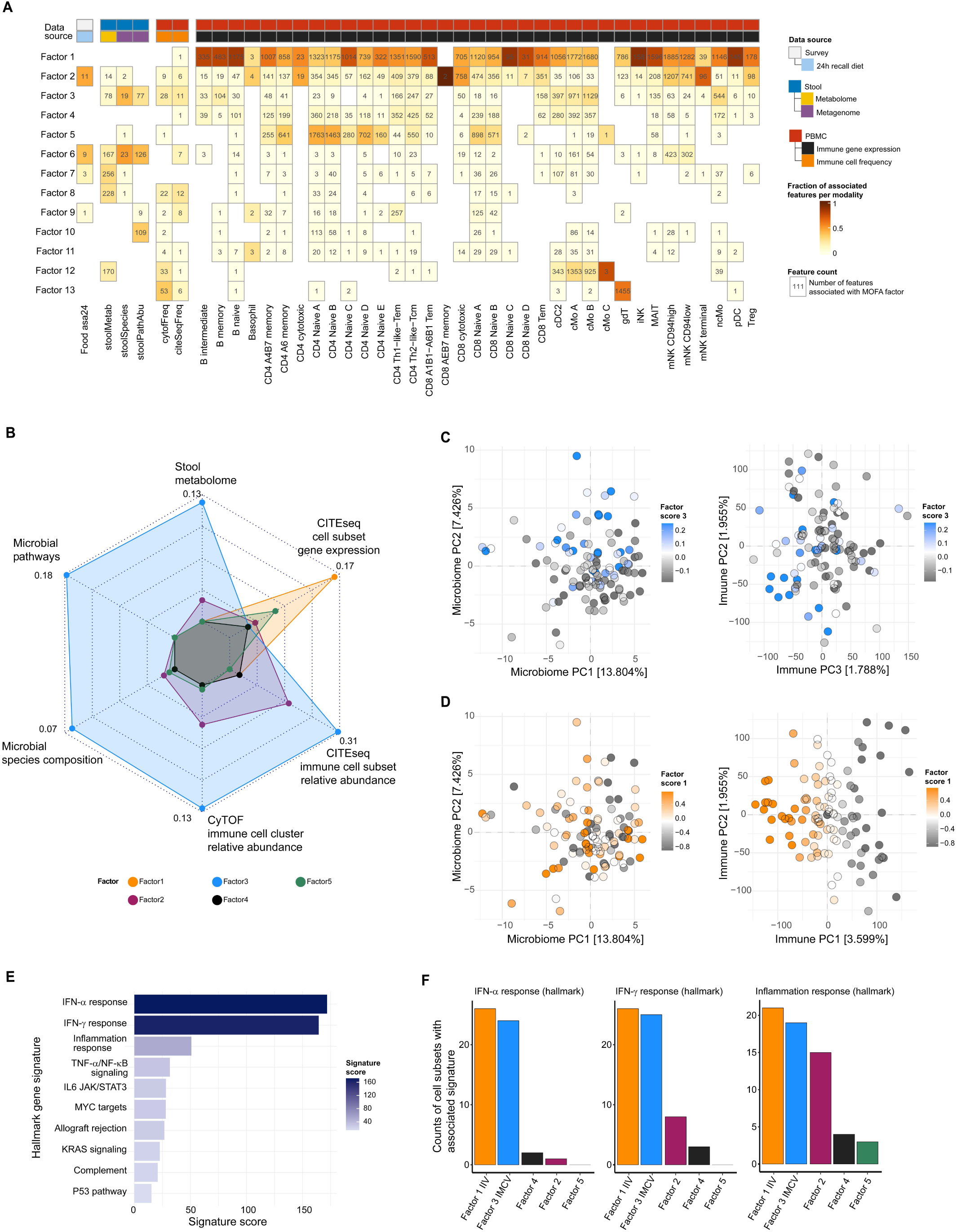
Integrative multi-omic analysis identifies microbiome-associated immune variation in interferon and inflammation responses. A. Magnitude of multi-omics features captured by MOFA factors. Labels show number of features significantly associated with each factor (adj.p < 0.1 and |feature weight| > 0.1). Colors quantify the proportion of features associated with each factor as a fraction of all significant features of the modality across MOFA factors. B. Variation captured by MOFA factors by modality. Values plotted are the fraction of features included in MOFA associated with each factor per modality. Values for CITE-Seq gene expression data are averages over all cell subsets. Each axis is scaled to the highest value (displayed). C-D. Distribution of study subjects obtained by principal component analysis of prevalent microbiome species composition data (C) and on CITE-Seq gene expression data (D). E. Combined transcriptional signature enrichment in IMCV (sum of -log10(adj.p)) across cell subsets from the CITE-Seq data. The top 10 dominant hallmark gene signatures are shown, ordered by signature score (sum of -log10(adj.p) across cell subsets). Signatures were evaluated using mHG test and hallmark gene sets. F. Number of cell subsets with significant enrichment of hallmark transcriptional signatures IFN-α response, IFN-γ response, inflammation response for the top 5 factors capturing the most variance across datasets. Signatures were evaluated using mHG test and hallmark gene sets.

To understand immune variation captured by IMCV, we first quantified the dominant signatures in the transcriptional data across factors. IFN alpha (IFN-α) response, IFN gamma (IFN-γ) response and inflammation response showed the highest signature scores (**Fig. 2E**). We then identified the top hallmark signatures for each factor and quantified the number of cell subsets that showed enrichment (**Fig. S2D**). IFN-α, IFN-γ, and inflammation response signatures were enriched in IMCV across the majority of immune cell populations (**Fig. 2F, S2D**). Interestingly, these signatures were also highly associated with IIV, which also captured additional signatures (**Fig. 2F, S2D**). This result further suggests that interferon and inflammation responses are dominant sources of transcriptional variability across the cohort, captured by IMCV and IIV.

### IMCV captures differences in the tonic-interferon response across the cohort

While IFN-related signatures were broadly captured by both IMCV and IIV (**Fig. 2F**), we hypothesized that IMCV and IIV might capture different elements of IFN response. Using IFN gene sets that do not overlap with inflammation-related signatures, we observed that most cell subsets showed positive enrichment for IFN signatures for both factors, though associations with IMCV were stronger for most immune cell populations (**Fig. 3A**). We next examined the specific genes that drive these signatures by evaluating differential gene expression across cell subsets in subgroups of individuals with the highest and lowest (20%) scores for each factor (**Fig. S3A**). The most variable and significant genes upregulated in the high IIV subgroups were largely expressed by lymphoid subsets (**Fig. 3B-C**, **Table S2**), while the most upregulated genes in the high IMCV group were enriched in myeloid cell subsets (**Fig. 3B**, **3D**). Most differentially expressed genes were unique to one factor or the other (**Fig. 3E**). Similar results were found when investigating the IFN-α response signature (**Table S2**).

**Figure 3.**
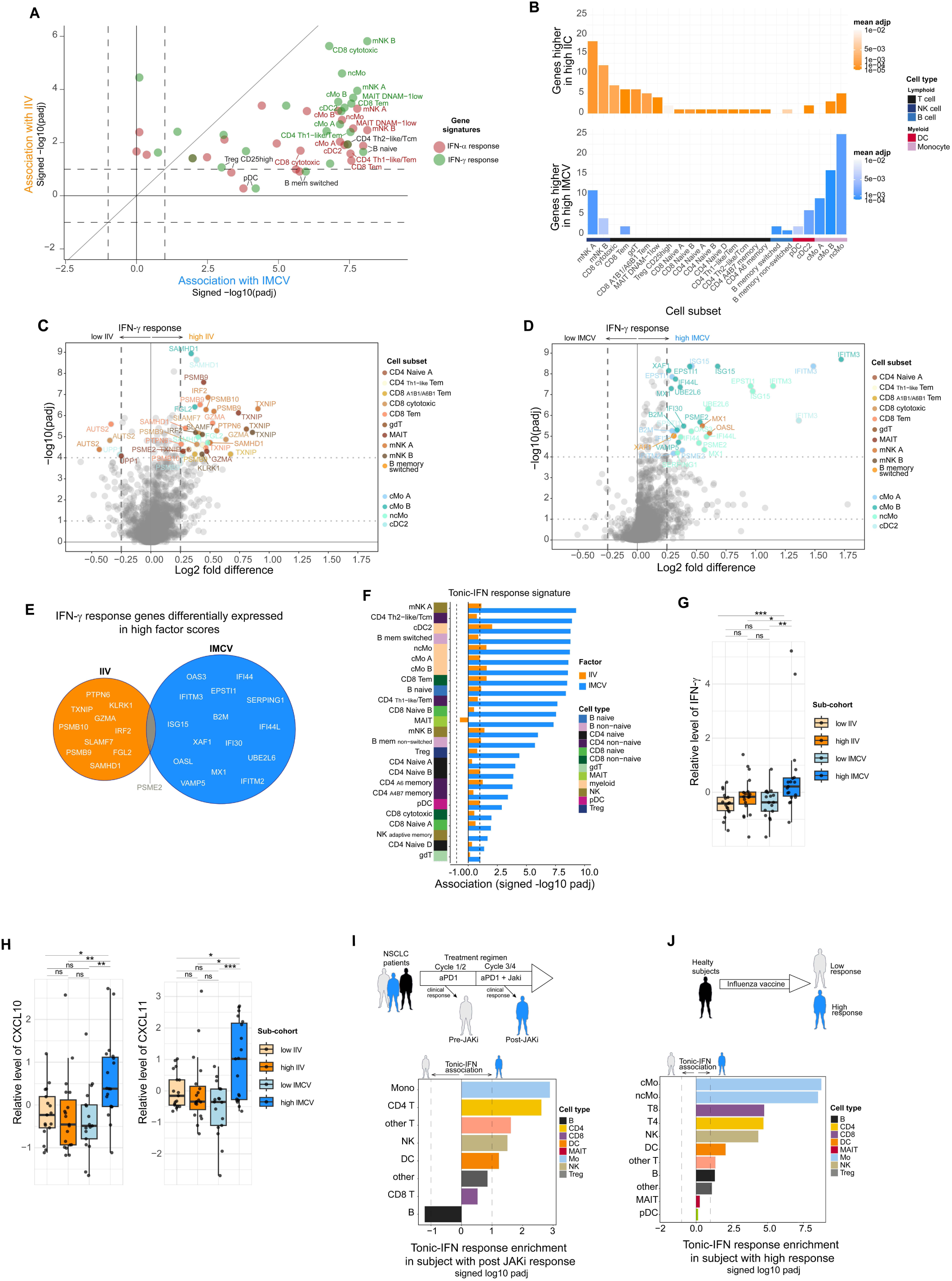
IMCV captures differences in the tonic-interferon response across the cohort. A. Comparison of the association of IIV and IMCV with IFN response signatures across cell subsets. mHG was used to test enrichment on CITE-seq gene expression data. Signed -log10(adj.p) for each factor is shown. B-E. Number of genes (B) and differential expression of IFN-γ response genes between study participants with high (top 20%) and low (bottom 20%) IIV (C) and IMCV (D) factor scores. Wilcoxon rank-sum test and BH correction were used. The most differentially expressed genes (adj.p <10^-4^ and |log2 fold difference| > 0.25) are considered and visualized on a Venn diagram (E). F. Association of IIV and IMCV with tonic-IFN response transcriptional signature across cell subsets. mHG enrichment test on CITE-seq gene expression data was used. Signed -log10(adj.p) for each factor is shown. G-H. Relative plasma levels of IFN-γ (G) and chemokines CXCL10 and CXCL11 (H) in sub-cohorts of ImmunoMicrobiome participants with high and low IIV and IMCV factor scores, with no overlap between the sub-cohorts. Kruskal-Wallis and Dunn post-hoc statistical tests were performed. ***=p<0.0005, **=p<0.005,*=p<0.05. I. Enrichment of the tonic-IFN gene signature in immune cell subsets of non-small cell lung cancer patients who responded to anti-PD1 alone or only responded after the combination with a JAK inhibitor. J. Enrichment of the tonic-IFN gene signature in immune cell subsets of individuals who mounted high or low responses to the seasonal *influenza* vaccine.

Interferon signaling induces different transcriptional responses depending on the duration and strength of the stimulus.^32^ Because of the differences in IFN response signatures between IMCV and IIV, we investigated their overlap with more nuanced and specific IFN response gene signatures. Genes strongly associated with IMCV were enriched for a “tonic-IFN response” gene signature, derived from an *in vivo* study that identified genes sensitive to steady-state, constitutively-expressed, low levels of IFN (**Fig. 3F, S3C**).^27^ They were also enriched for a related “prolonged-IFN response” signature, composed of ISGs expressed in response to prolonged low doses of IFN in human cells *in vitro* (**Fig. S3B-D**),^33^ suggesting potential differences in IFN exposure across the cohort. In contrast, associations between IIV and these signatures were much weaker across cell types (**Fig. 3E, S3B**).

We therefore hypothesized that circulating IFN levels might be higher in the high-IMCV subgroup of study participants. We measured cytokine levels in plasma specimens and found higher levels of IFN-γ, along with CXCL10 and CXCL11 chemokines encoded by ISGs, in high-IMCV study participants (**Fig. 3G-H**). Moreover, plasma levels of these molecules also correlated with tonic-IFN response and prolonged-IFN response gene signatures across cell subsets (**Fig. S3E-F**).

We hypothesized that interindividual variation in tonic-IFN states could reflect steady-state differences that impact responses to immunomodulatory treatments. Therefore, we analyzed transcriptional data from an *influenza* vaccine study.^34^ We observed that the baseline immune states of individuals who went on to mount strong immune responses to the vaccine were associated with the same tonic-IFN response signature (**Fig. 3I**). Notably, recent history of vaccination in the ImmunoMicrobiome cohort showed only a modest association between influenza vaccine status and IIV factor score and no association with IMCV (**Fig. S3G**). Conversely, analysis of patients with non-small cell lung cancer treated with immunotherapy^35^ revealed that the tonic-IFN response signature was associated with patients who only responded to anti-PD1 checkpoint blockade after the addition of the JAK inhibitor itacitinib, which blocks IFN signaling (**Fig. 3J**).

Taken together, these data show that IIV and IMCV are associated with distinct IFN-related transcriptional programs. Elevated expression of a tonic-IFN signature in IMCV, especially prominent in myeloid cell subsets, coincides with higher response to *influenza* vaccination and is elevated in cancer patients who benefit from combined JAK inhibition with anti-PD1.

### Inflammation and TGF-β transcriptional programs are differentially regulated between IMCV and IIV

Our prior analysis indicated that inflammation-related gene signatures were also among the most variable pathways across the cohort (**Fig. 2E**) and were associated with IMCV and IIV (**Fig. 2F, S2D**). In fact, IIV and IMCV factor scores were positively associated with IFN signatures and negatively associated with inflammation-related signatures (**Fig. 4A**). In the 3 major monocyte clusters identified by CITE-seq, both IMCV and IIV exhibited negative associations with inflammation response and TNF-α/NF-κB signaling signatures (**Fig. 4B**). Despite similar overall relationships with these gene signatures, these were driven by a distinct set of differentially expressed genes for each factor (**Fig. 4C-D, Table S3**). Key IL-1 pathway genes were expressed at lower levels in individuals with high IMCV scores, including the cytokines *IL1A* and *IL1B*. In contrast, the high-IIV group exhibited lower expression of IL-6 regulators, including oncostatin M (OSM)^36^ and *TIMP1*.^37^ Similar results were observed with the TNF-α/NF-κB signaling signature (**Fig. 4D, Table S3**), which additionally highlighted higher levels of TLR/IFN-induced transcription factor *ATF3* that represses NF-KB inflammatory signaling^38^ in the high-IMCV group. Lipopolysaccharide-induced TNF-α factor (*LITAF*) that induces TNF-α expression^39^ was expressed at higher levels in the high-IMCV group but lower in the high-IIV group (**Fig. 4D**). In sum, these results indicate that both IMCV and IIV are inversely associated with inflammation and TNF pathways in monocytes but through distinct sets of genes (**Fig. S4A**).

**Figure 4.**
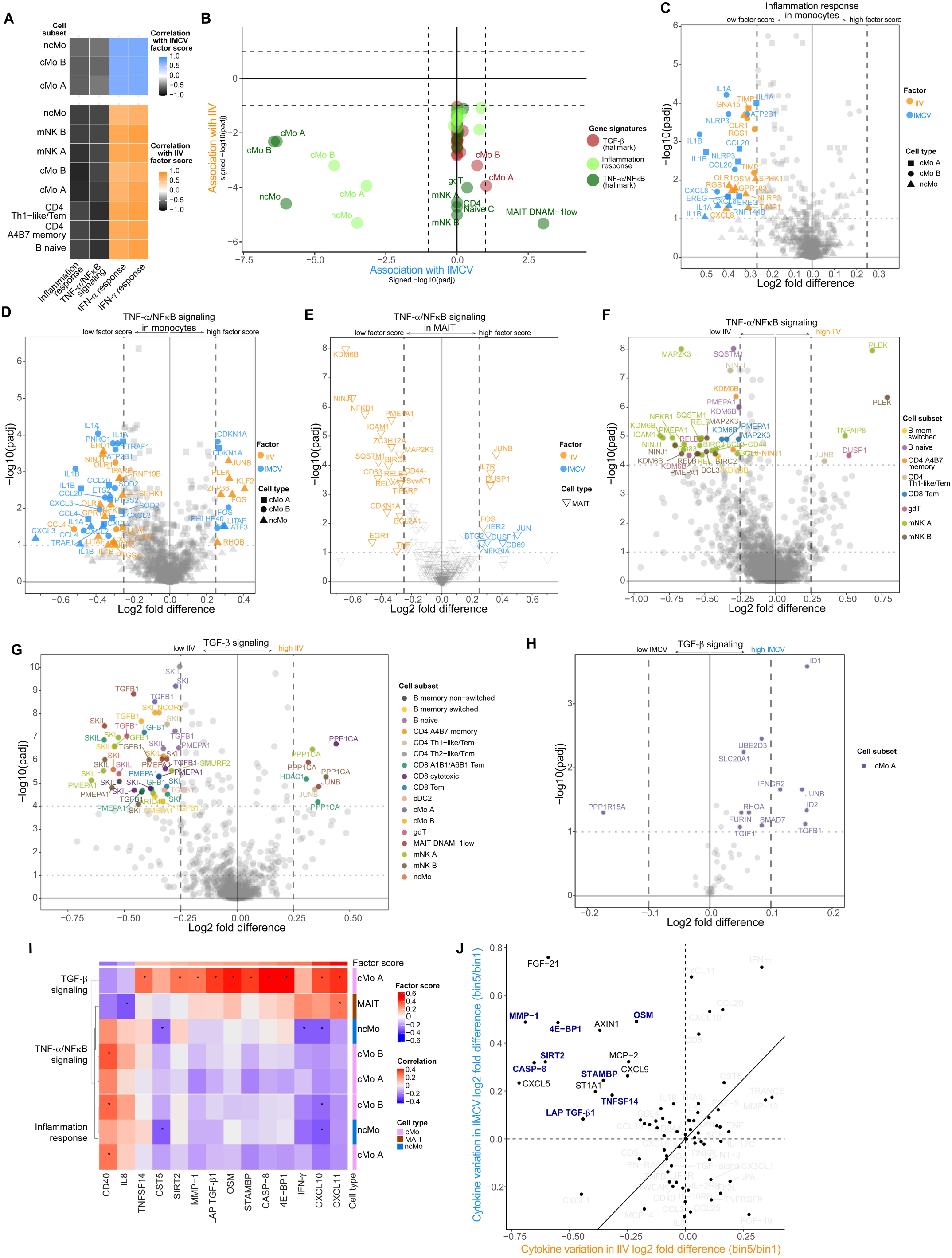
Inflammation and TGF-β transcriptional programs are differentially regulated between IMCV and IIV. A. Correlation of IIV and IMCV factor scores with IFN and inflammation-associated signatures (sum of weighted expression of core genes). B. Association of IIV and IMCV with inflammation-associated signatures across cell subsets. mHG was used to test enrichment on CITE-seq gene expression data. Signed -log10(adj.p) for each factor is shown. C-E. Inflammation-related genes differentially expressed between study participants with high and low factor scores. Inflammation response genes (C) or TNF-α/NF-κB signaling genes (D) in monocyte subsets and TNF-α/NF-κB signaling genes in MAIT cells (E) are shown. Wilcoxon rank-sum test and BH correction were used. Most differentially expressed genes (adj.p <10^-1^ and |log2 fold difference| > 0.25) are annotated and colored by factor. F. TNF-α/NF-κB signaling genes differentially expressed between study participants with high and low IIV factor scores. Wilcoxon rank-sum test and BH correction for multiple testing were used. Most differentially expressed genes (adj.p <10^-4^ and |log2 fold difference| > 0.25) are annotated and colored by cell subset. G-H. Same as F for TGF-β signaling signature in high and low IIV factor scores (G) or IMCV factor scores (H). I. Plasma cytokine association with IMCV and inflammation-related signatures across cell subsets. Spearman rank correlation was used to evaluate the associations. |r| > 0.25 indicated with an asterisk. J. Plasma cytokine variation between high and low factor scores in IIV and IMCV respectively. Cytokines that are simultaneously higher in the IMCV high cohort and lower in the IIV high cohort are annotated and cytokines associated with cMo-A TGF-β signatures are colored.

In many other cell subsets, IIV was negatively associated with inflammation-related signatures, while IMCV had no association (**Fig. 4B**). MAIT cells, a subset of invariant T cells recognizing vitamin metabolites, including those derived from the gut microbiome,^40^ were the only subset in which the TNF-α/NF-κB signaling signature was positively associated with IMCV but negatively associated with IIV (**Fig. 4B)**. Key genes driving the negative association with IIV included both TNF itself as well as genes encoding NF-κB subunits and their regulators (**Fig. 4E, Table S3**). Conversely, the positive association with IMCV was driven by higher expression of inflammation-associated factors downstream of TNF-α. Interestingly, the transcription factor *JUNB,* which acts as a negative regulator of JUN,^41^ was expressed at higher levels in the high-IIV subgroup, suggesting differences in the AP-1 pathway regulation in MAIT cells. While there was no association between TNF-α/NF-κB signaling gene expression and IMCV in other lymphocyte subsets, the genes driving negative associations with IIV largely overlapped with those observed in MAIT cells (**Fig. 4F**).

We also investigated associations between these factors and the TGFβ pathway, which was uniquely negatively associated with IIV (**Fig. 4B**). Individuals in the high-IIV group exhibited lower TGFB1 expression across multiple cell subsets and higher level of TGF-β receptor inhibitor *PPP1CA* (**Fig. 4G**, **Table S3**). In contrast to IIV, the TGF-β signature in cMo-A was positively associated with IMCV (**Fig. 4B**), driven by modestly higher expression of *TGFB1*, *IFNGR2*, *JUNB*, transcription factors *ID1* and *ID2,* and lower expression of *PPP1R15A*, involved in *TGFBR1* inhibition (**Fig. 4H**).^42^

We further evaluated circulating factors associated with these pathways. Both inflammation and TNF-α/NF-κB signaling signatures in monocytes were positively associated with circulating levels of CD40, which signals through the NF-κB pathway (**Fig. 4I**). TGF-β signature in cMo-A correlated with plasma levels of the latent complex LAP-TGF-β1, as well as other factors associated with IMCV, including ISGs CXCL10 and CXCL11 (**Fig. 4I**). Consistent with transcriptional data, soluble LAP-TGF-β1 levels were negatively associated with IIV (**Fig. 4J**). Taken together, these data show that high-IMCV and high-IIV subgroups exhibit key differences in inflammation response and TGF-β signaling pathways.

### IMCV captures differences in SIGLEC-1^high^ monocytes and PD1^high^ICOS^high^ memory T cells across the cohort

IMCV was further distinguished from the IIV by its association with immune cell population abundances (**Fig. 2A**). The relative abundance of monocyte cell subsets that exhibited distinct interferon and inflammation response transcriptional signatures was positively associated with IMCV (**Fig. S5A**). The CyTOF data, which allowed for deeper sampling due to its higher throughput, confirmed that the relative abundances of several monocyte clusters were positively associated with IMCV (**Fig. 5A**). When compared to non-associated clusters, these monocytes exhibited higher protein expression of the MHC class II protein HLA-DR and the Fc gamma receptor CD64, which are induced by IFN (**Fig. 5B-C**).^43–46^ In addition, these cells expressed higher levels of myeloid activation markers (CCR7, CD14), as well as the scavenger receptor CD163, associated with an anti-inflammatory phenotype (**Fig. 5B-C**).^47^ Consistent with these results, CITE-Seq classical monocyte cluster B (cMo-B), the abundance of which was associated with IMCV, also displayed higher expression of MHC class II pathway genes (HLA-DP/DQ/DR and CD74), when compared to other cMo subsets. Similarly, we found higher expression of the immunoproteasome gene *PSME2*, and calreticulin (*CALR*), a chaperone critical for antigen processing and presentation, along with several ISGs (**Fig. 5D**). Conversely, cMo-B had lower expression of pro-inflammatory genes, including AP-1 subunits (*JUN* and *FOS*), and calprotectin (*S100A8*/*S100A9*), an endogenous ligand of TLR4 (**Fig. 5D**). To further investigate the differences of phenotype of these myeloid cell subsets captured by IMCV, we compared CITE-seq protein levels in individuals with high and low IMCV factor scores. SIGLEC-1 (CD169), a marker previously shown to define IFN-induced monocytes with enhanced antigen presentation capability,^48^ protection against sepsis in mice^49^ or severe COVID in humans,^50^ was more highly expressed by monocytes in high-IMCV individuals (**Fig. 5E, S5B**). Although cMo-B expressed the highest level of SIGLEC-1, other myeloid subsets also showed similar differential expression in high-IMCV study participants, suggesting that elevated SIGLEC-1 expression may be characteristic of a broader myeloid state in these individuals (**Fig. S5B**).

**Figure 5.**
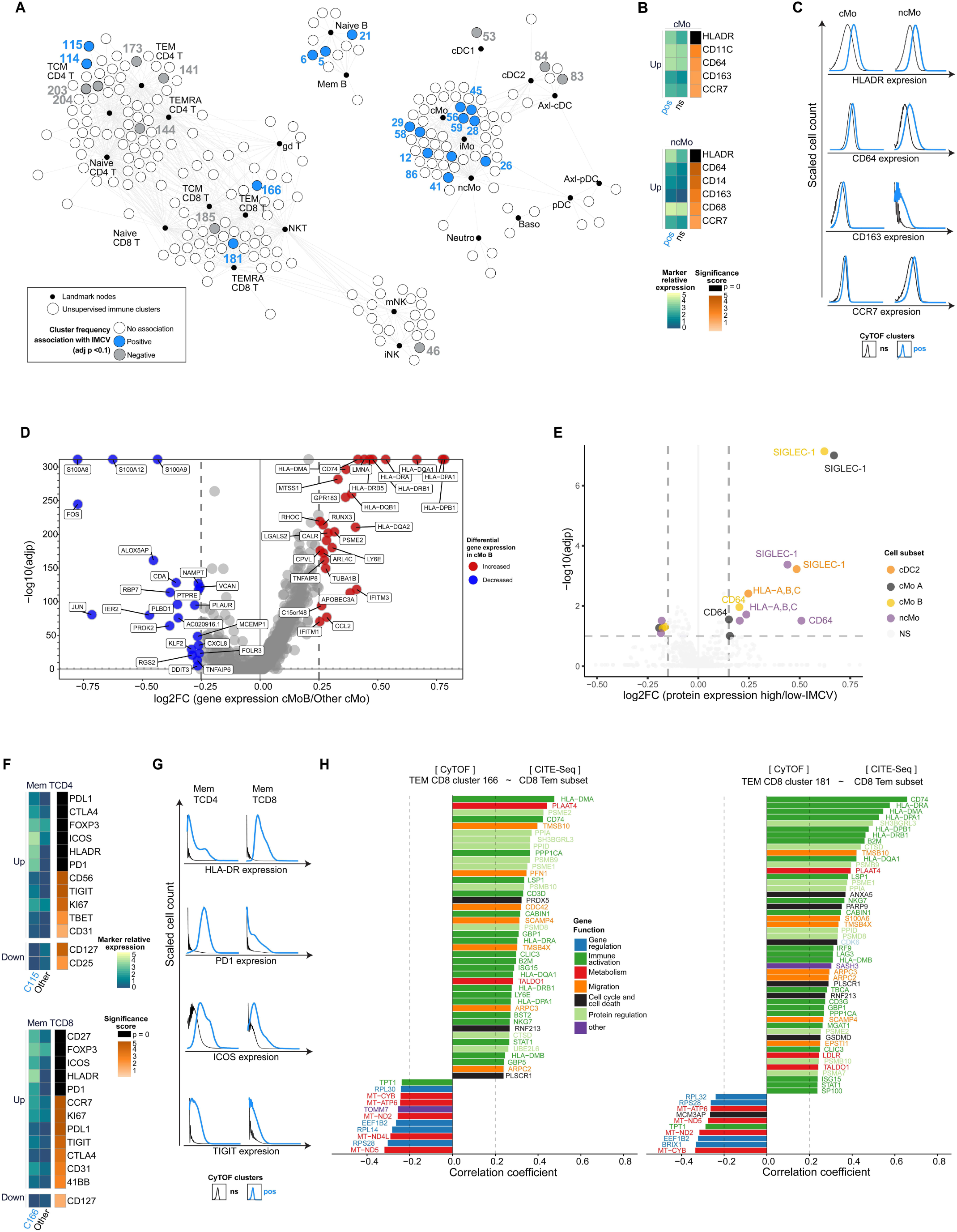
IMCV captures differences in SIGLEC-1^high^ monocytes and PD1^high^ICOS^high^ memory T cells across the cohort. A. SCAFFoLD map of CyTOF immune cluster frequency associated with IMCV. Unsupervised immune clusters (white and colored nodes) are mapped onto reference immune populations (black nodes). Linear regression and BH correction (adj.p < 0.1) was used to evaluate association of features with IMCV B. Proteins differentially expressed in combined CyTOF monocytes clusters whose frequencies are positively associated (Pos) and not significantly associated (ns) with IMCV. Kruskal–Wallis and post-hoc pairwise Wilcoxon tests were used to evaluate proteins differentially expressed (p<0.00001). Scaled medians of expression and significance are shown. C. Expression of selected surface proteins shown in B. D. Differential CITE-Seq gene expression between cMo-B and other classical monocyte cell subsets in CITE-seq data. Features with adj.p < 0.1 and |log2 fold-change| > 0.25 are colored and annotated. E. Differential CITE-Seq protein expression between study participants with low and high IMCV factor scores. Features with adj.p < 0.1 and |log2 fold-change| > 0.2 are colored and annotated. F. Same as B, with memory CD4 T cell cluster 115 and CD8 T cell cluster 166. G. Same as C, with combined memory CD4 T cell clusters 114/115 and CD8 T cell clusters 166/181. H. Correlation between CITE-seq gene expression in CD8 effector memory T cell subset and CyTOF frequency of CD8 effector memory T cell clusters C166 and C181. Genes with correlation coefficient |r| > 0.2 are displayed and annotated. Coloring indicates manually curated broad functions (see table S4).

Several memory T cell clusters were also positively associated with IMCV (**Fig. 5A**). Memory CD4 T cell cluster 115 (C115) and CD8 T cluster 166 (C166) exhibited higher expression of a combination of costimulatory and coinhibitory checkpoint molecules compared to clusters that were not associated with IMCV (**Fig. 5F**). These clusters also expressed higher levels of the activation marker HLA-DR and transcription factor Foxp3 and lower expression of cytokine receptors. Two additional clusters positively associated with IMCV displayed similar phenotypes except for the expression of Foxp3 (**Fig. 5G**, **S5C**). When compared to Tregs (**Fig. S5D-E**) and memory T cells not associated with IMCV (**Fig. S5F-G**), combined single cells from the four clusters positively associated with IMCV displayed higher expression of Foxp3 and most checkpoint molecules, including inhibitory checkpoint ligand PD-L1. CD4 and CD8 memory T cell clusters that were negatively associated with IMCV (**Fig. 5A and S5A**) generally revealed the opposite expression profile (**Fig. S5D-G**). Finally, to further examine the phenotype of the CD8 TEM cells associated with IMCV, we identified correlations between cluster abundances in the CyTOF data and gene expression in the CITE-seq data for this cell population. Most positively correlated genes were related to T cell activation, including many antigen processing and presentation molecules (**Fig. 5H, Table S4**), consistent with the results from CyTOF. Together, these results suggest that IMCV is associated with memory T cells with an activated phenotype and elevated expression of checkpoint molecules.

### Established immunomodulatory gut microbiome pathways and molecules are associated with IMCV

The most unique feature of IMCV was its coordination between immune and microbiome datasets (**Fig. 2A**). The microbiome features associated with IMCV included 77 microbial pathways (**Table S5**), including several related to production of known immunomodulatory molecules: biosynthesis of short-chain fatty acids (SCFA), metabolism of polyamines, and production or modification of bacterial cell wall components lipopolysaccharide (LPS) and peptidoglycans (**Fig. 6A**).^51–54^ In addition, 78 stool metabolites were also associated with IMCV (**Table S5**), including many of the immunomodulatory molecules produced by these pathways (**Fig. 6B**). The SCFAs acetate, butyrate, and propionate were positively associated with IMCV, as were SCFA-related compounds. Inositol, a precursor of propionate, was positively associated with IMCV, while its phosphorylated precursor, inositol-P,^55^ was negatively associated. Out of 8 polyamines detected in stool, 6 were positively associated with IMCV, including ornithine that had precursors negatively associated with IMCV (**Fig. S6A**). Primary bile acids chenodeoxycholic acid and cholic acid, also studied for their immunomodulatory effects,^56^ were associated with IMCV as well. Together, these findings reveal that the immune states captured by IMCV are coincident with immunomodulatory pathways in gut microbiota and the metabolites they produce.

**Figure 6.**
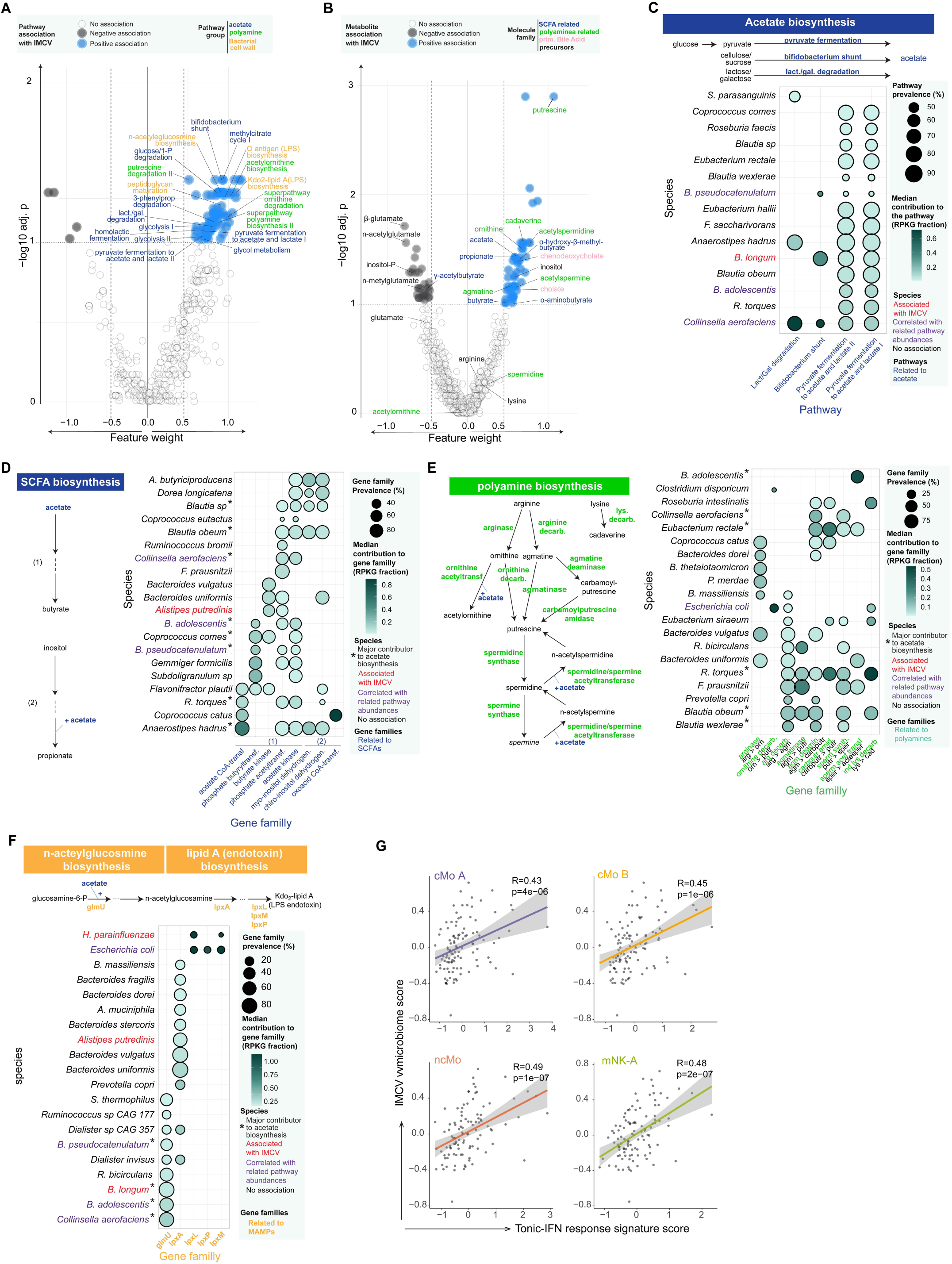
Established immunomodulatory gut microbiome pathways and molecules are associated with IMCV. A. Microbiome pathway abundances association with IMCV. Linear regression and BH correction for multiple testing (adjp < 0.1) and and MOFA feature weight(|FW| > 0.45) were used to determine association of features with IMCV. Pathways related to SCFA, bacterial cell wall, and polyamines are annotated. B. Stool metabolite association with IMCV. Linear regression and BH correction for multiple testing (adjp < 0.1) and and MOFA feature weight(|FW| > 0.45) were used to determine association of features with IMCV. SCFA, polyamines, primary bile acids and their precursors are annotated. C. Species contribution to acetate biosynthesis pathway total genomic content. Major contributors are shown (prevalence >25% and a median contribution > 0.05). All contributions, including from minor and unclassified species, are documented in Table S3. D-F. Species contribution to gene families total genomic content for SCFA biosynthesis (D), polyamine metabolism (E), Kdo2-lipid A (LPS-endotoxin) biosynthesis (F). Major contributors are shown (top 20 most prevalent species, median contribution > 0.05). All contributions, including from minor and unclassified species are documented in the Table S3. G. Microbiome composite score (median of microbiome features associated with IMCV) relationship with tonic-IFN response gene signature (median expression of core genes). Spearman rank correlation was used to determine coefficients of correlation and significance. Linear regression lines are shown. Cell subsets with the highest correlations are shown.

Next, we investigated the bacterial species encoding these pathways associated with IMCV. We measured their contribution to pathway abundances through the prevalence and average copy number of the relevant genes they carried across the cohort (**Table S6**). *Bifidobacterium longum*, *Bifidobacterium adolescentis*, *Bifidobacterium pseudocatenulatum*, and *Collinsella aerofaciens* were among the major contributors to acetate biosynthesis pathway abundance (**Fig. 6C**). The relative abundance of these species was also correlated with various acetate biosynthesis pathway abundances (**Fig. S6B**). *B. longum* abundance was directly associated with IMCV (**Fig. S6C**). We also investigated a non-exhaustive set of metabolic reactions that contribute to the transformation of acetate to butyrate (**Fig. S6D**) and inositol to propionate and found that several of the major contributors were the same species involved in acetate biosynthesis (**Fig. 6D**). Of these, *Blautia obeum* and *R. torques* also encoded the alkaline phosphatase gene family involved in catalyzing the dephosphorylation of inositol-P, suggesting a potential role for these bacteria in propionate biosynthesis (**Fig. S6E**).

A partially-overlapping group of species contributed to pathway abundances related to the metabolism of polyamines (**Fig. 6E**). Several species had genes encoding the transformation of agmatine to putrescine with high prevalence. The same species were also major contributors to the subsequent production of spermidine and spermine. *B. adolescentis* was the highest contributor in gene family abundance to the acetylation of spermine/spermidine, while *R. torques* and *B. obeum* were important contributors of gene family content for cadaverine biosynthesis from lysine. Overall, the complementarity of species contributing to polyamine metabolism was suggestive of community metabolism. Consistent with this notion, the abundance of these individual species was not associated with IMCV (**Fig. S6C**) nor correlated to polyamine pathway unstratified abundances (**Fig. S6F**), highlighting the importance of examining metabolic pathways and metabolite levels in addition to microbiota composition.

Examining the species that contributed to biosynthesis pathways of LPS, which is detected by the innate immune receptor TLR4, revealed further overlap. *Bifid*o*bacterium* species and *C. aerofaciens* were also important contributors to the biosynthesis of n-acetylglucosamine (**Fig. 6F**), a precursor of LPS. The abundance of these species further correlated with LPS biosynthesis pathway abundances (**Fig. S6G**). The subsequent transformation of n-acetylglucosamine into LPS endotoxin KDo2-lipid A associated with IMCV and was dominated by distinct species, including *E. coli* and *Haemophilus parainfluenzae*, that uniquely carried genes involved in the final steps of its biosynthesis. Several major contributors to these pathways were positively associated with IMCV (**Fig. S6C**), and several overlapped with those for SCFA and polyamine metabolism pathways. Some of these species were also major contributors to the abundance of the bile salt hydrolase gene family that can produce the unconjugated bile acids associated with IMCV (**Fig. S6H**).

We next examined the phylogenetic relationship of species carrying genes involved in the production of molecules associated with IMCV (SCFA, polyamines, LPS endotoxin, primary bile acids). These species were phylogenetically diverse, spanning more than 14 families. A core community of species, which encoded genes involved in all key processes associated with IMCV, spanned 5 microbial families (**Fig. S6I**). Of note, the metabolic pathways encoded by these microbes and metabolite levels were generally more strongly associated with IMCV than were the abundances of bacterial species. Altogether, these results highlight a complex collection of phylogenetically diverse microbial species defined by their genetic content encoding pathways involved in the production of immunomodulatory molecules captured by IMCV.

Lastly, we determined whether the IFN pathways associated with IMCV were directly associated with these microbiome features. As expected, a IMCV microbiome composite score of key microbiome pathway abundances and stool metabolites was significantly correlated with the tonic-IFN response signature across the immune cell subsets with the highest IFN signature scores, including monocytes and a subset of mature NK cells (cluster mNK-A)(**Fig. 6G**).

### Individuals with a microbiome-associated high-tonic interferon immune state exhibit increased SIGLEC-1^high^ monocytes and CD69^high^ innate lymphoid immune subsets

While IIV and IMCV capture unique immune variability across the full cohort of healthy participants, participants can be specifically classified based on their factor scores to identify distinct immune states (**Fig. 7A**). We defined sub-cohorts of the top 20 individuals with the highest scores for each factor (**Fig. S7A**). The sub-cohorts were distinct except for two participants among the top 20 for both factors, whom we excluded from subsequent analysis. Differential gene expression analysis between high-IMCV and high-IIV participants followed by enrichment analysis revealed a relative enrichment of signatures for IFN-α and IFN-γ response, as well as tonic and prolonged-IFN response signatures, across myeloid cell subsets and several innate lymphoid cell subsets in the high-IMCV sub-cohort (**Fig. 7B**). Consistent with the system-wide negative association of IIV with inflammation response, TNF-α/NF-κB signaling, and TGF-β signaling signatures (**Fig. 4B**), these were lower in the high-IIV sub-cohort across a variety of cell subsets (**Fig. 7B**). However, the inflammation signature in cMo-A and cMo-B was higher in the high-IIV sub-cohort (**Fig. 7B**).

**Figure 7.**
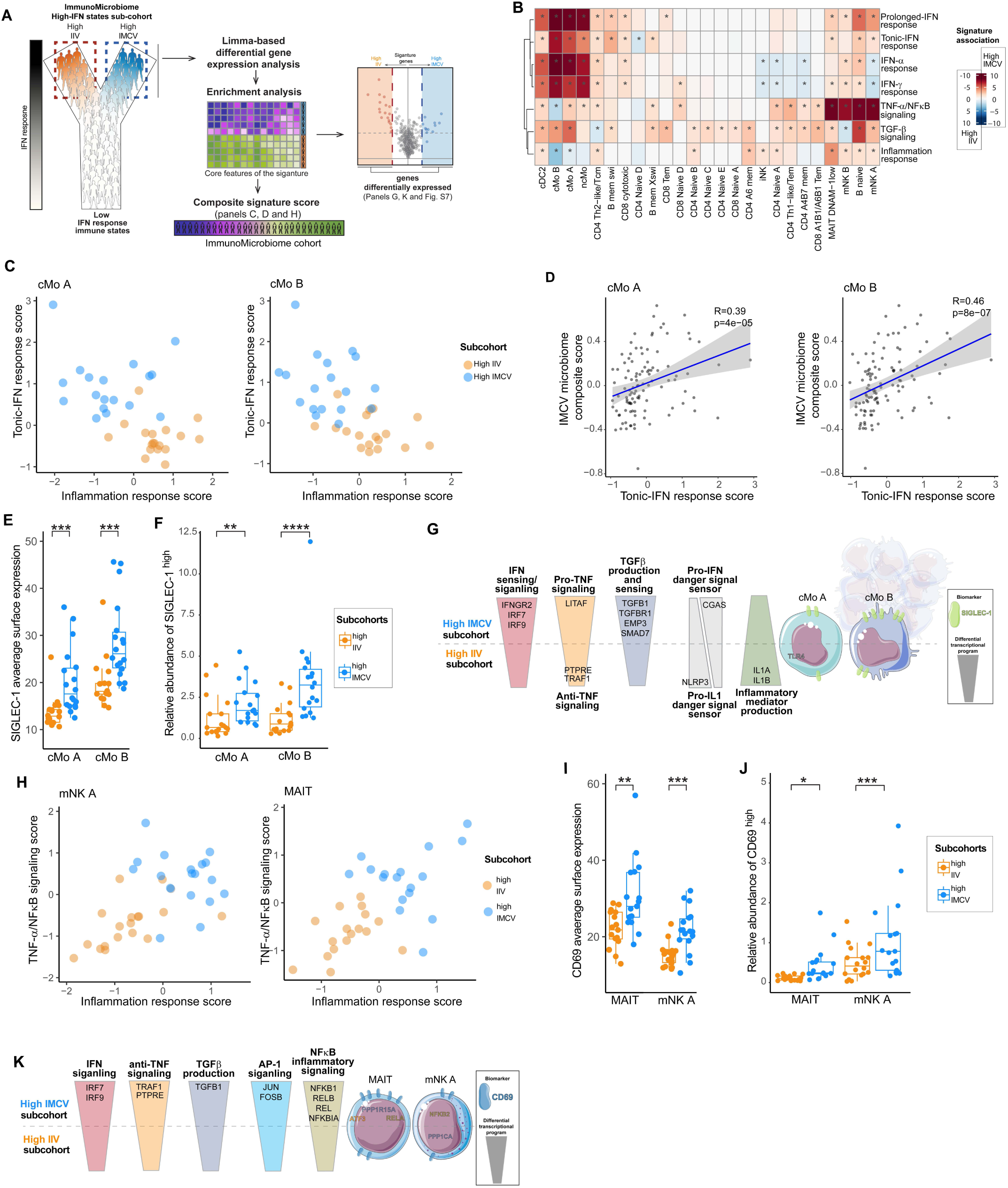
Individuals with a microbiome-associated high-tonic interferon immune state exhibit increased SIGLEC-1^high^ monocytes and CD69^high^ innate lymphoid immune subsets. A. Schematic of the cohort and the analytical approach performed in Fig.7. Genes differentially expressed between high-IMCV and high-IIV sub-cohorts were identified using limma, ranked using limma’s log2 fold-difference and used for enrichment analysis for transcriptional signatures of interest. Core genes driving the signatures are used to build individual composite signature scores (median expression of core genes), which were used to map study participants based on their immune states (panels C,H). For enriched signatures, differential gene expression between high-IMCV and high-IIV sub-cohorts are evaluated post hoc using Wilcoxon rank-sum test (Panels G, K and Fig. S7 B-O). B. Association of high-IMCV and high-IIV with transcriptional signatures across cell subsets from CITE-seq gene expression data. Associations are tested using mHG enrichment analysis. Significance (adj.p < 0.1) is indicated by an asterisk symbol. Color scale represents signed -log10 (adj.p). C. Scatter plot mapping study participants from the high-IMCV and high-IIV sub-cohorts. Individual coordinates derived from their composite signature scores (median expression of core genes) for tonic-IFN response and inflammation response in monocyte subsets cMo-A and cMo-B are used. D. Relationship of study participants’ microbiome composite score and Tonic-IFN response transcriptional signature score. The microbiome composite scores were derived from microbiome features associated with IMCV. Spearman rank correlation was used to determine coefficient of correlation and significance (shown) and linear regression lines are used. E-F. Comparing cell surface expression of activation biomarker SIGLEC-1 (E) and relative abundance of SIGLEC-1^high^ (F) cMo-A and cMo-B subsets of study participants from high-IMCV and high-IIV sub-cohorts in CITE-Seq data. G. Schematic summarizing key differential gene expression and surface biomarker between high-IIV and high-IMCV sub-cohorts in cMo-A and cMo-B (see Fig. S7B-H). H. Scatter plot mapping study participants from the high-IMCV and high-IIV sub-cohorts. Unique coordinates derived from their composite signature scores for TNF/NF-κB and Inflammation in mNK A and MAIT subsets are used. I-J. Comparing cell surface expression of activation biomarker CD69 (I) relative abundance of CD69^high^ (J) MAIT and mNK A subsets in study participants from high-IMCV and high-IIV sub-cohorts in CITE-Seq data. K. Schematic summarizing key differential gene expression and surface biomarker between high-IIV and high-IMCV sub-cohorts in MAIT and mNK A subsets (see Fig. S7I-O).

While these gene signatures were derived from the literature, we explored whether individual composite scores, based on the expression of the core genes driving the enrichment of each signature, might further summarize immunological states in these sub-cohorts (**Table S7**). Indeed, mapping individuals based on data-driven tonic-IFN response and inflammation response composite signature scores in monocytes distinguished the sub-cohorts from one another and displayed the opposing nature of these transcriptional programs in healthy individuals (**Fig. 7C**). The tonic-IFN response composite score in cMo-A and cMo-B was significantly associated with the microbiome score made of features associated with IMCV (**Fig. 7D**). Among the genes in this signature, IFNGR2, IRF7 and IRF9 involved in IFN-γ signaling were higher in the high-IMCV cohort compared to the high-IIV cohort (**Fig. S7B-C**), consistent with their higher IFN response signatures and expression of ISGs (**Fig. S7B**, **S7D**). Inflammation-related genes were also differentially expressed between the sub-cohorts, as expected. Monocytes in the high-IMCV cohort expressed higher levels of TNF-α signaling molecule LITAF and lower levels of regulators of TNF-α signaling (**Fig. S7E**), along with higher expression of TGF-β and elements of TGF-β sensing/signaling machinery (**Fig. S7F**). Expression of innate immune pattern recognition receptors was also different; monocytes in the high-IMCV sub-cohort exhibited higher expression of cytosolic DNA sensor CGAS, which drives IFN production, and lower expression of NLRP3 (**Fig. S7G**), which promotes IL-1 maturation, as well as IL1A and IL1B (**Fig. S7H**).

Consistent with our observation that SIGLEC-1 was expressed at a higher level in monocytes of individuals with high-IMCV factor scores (**Fig. 5E**), SIGLEC-1 was also expressed at a higher level in monocytes of participants in the high-IMCV sub-cohort compared to the high-IIV sub-cohort (**Fig. 7E**), resulting in a higher relative abundance of SIGLEC-1^high^ monocytes (**Fig. 7F**). Together, these data highlight that the immune system of individuals with a high tonic-IFN immune state associated with microbiome features is characterized by SIGLEC-1^high^ monocytes with high IFN response and low inflammation transcriptional states (**Fig. 7G**).

MAIT and mNK-A lymphoid cell subsets also displayed differences between the high-IIV and -IMCV sub-cohorts (**Fig. 7B**). Like monocytes, these cell subsets expressed higher levels of tonic/prolonged-IFN ISGs and IFN signaling machinery in the high-IMCV sub-cohort (**Fig. S7I-J**), but they also showed higher enrichment for inflammation response and TNF-α/NF-κB signaling transcriptional gene signatures in the high-IMCV sub-cohort. The composite score of these signatures also stratified the high-IMCV and high-IIV sub-cohorts (**Fig. 7H**). These subsets in the high-IMCV sub-cohort expressed higher levels of NF-κB-related genes (**Fig. S7K**) as well as AP-1 subunits *JUN* and *FOSB* (**Fig. S7L**). In contrast to the monocytes, MAIT and mNK-A cells in participants from the high-IMCV sub-cohort expressed higher levels of regulators for TNF-α signaling (**Fig. S7M**). Both cell subsets also expressed higher levels of TGF-β and differential expression of TFGb pathway signaling molecules in participants from the high-IMCV sub-cohort (**Fig. S7N**).

Since microbial cues can activate these lymphoid cell subsets,^57^ we additionally examined expression of the activation marker CD69, which was expressed at higher levels in MAIT and mNK-A cell subsets in the high-IMCV cohort (**Fig. 7I** and **S7O**), resulting in a higher relative abundance of CD69^high^ cells (**Fig. 7J**). These data highlight that activated innate lymphoid cell subsets found in the high-IMCV sub-cohort can be identified by their unique transcriptional states and expression of the cell surface marker CD69 (**Fig. 7K**).

Altogether, our results show that individuals with high-IFN immune states can be classified based on IMCV and IIV factor scores and exhibit key immunological differences. The former harbor SIGLEC-1^high^ monocytes with elevated expression of tonic-IFN response genes, reduced expression of inflammatory mediators, and increased TGF-β signatures. In addition, their MAIT and mNK-A cell subsets exhibited increased expression of CD69, NF-κB and AP-1 signaling machinery, and genes involved in TGF-β signaling.

## Discussion

This study presents a comprehensive multi-omic dataset exploring variation in the microbiome and the immune system in healthy individuals. Using a factor-based integrative approach, we identified 2 axes of immune variation involving IFN response programs. One axis encompassed specific microbiome pathways, metabolites, and microbial communities that covaried with immune cell abundances and gene expression. Individuals displaying elevated levels of these features showed a distinct microbiome-associated immune state, marked by heightened tonic-IFN gene expression across immune cell types, expanded SIGLEC-1^high^ monocytes with downregulated inflammation transcriptional programs, as well as activated MAIT and NK cell states with increased CD69 expression.

Clinical studies have identified associations between the microbiome and immunological states in pathological settings.^15,50,58–60^ However, the extent to which the microbiome and immune system are coordinated in humans in the absence of pathology has been unclear. Among the immune features we identified to be coordinated with the microbiome in healthy individuals, several have been associated with clinical outcomes in patients. For example, elevated levels of HLA-DR^high^ SIGLEC-1^+^ monocytes in circulation at baseline were found to be protective of severe COVID disease, while expression of the alarmins S100A8 and S100A9 was elevated in patients with severe disease.^50^ This is consistent with our finding that SIGLEC-1^high^ monocytes associated with IMCV express lower levels of *S100A8* and *S100A9* transcripts compared to other monocyte subsets (**Fig. 5D**). Another study found that SIGLEC-1^high^ monocytes had heightened T cell activation capacity.^48^ This is consistent with our finding that SIGLEC-1^high^ monocytes associated with IMCV have higher levels of HLA-DR surface protein (**Fig. 5B-C**) and genes encoding MHC class II molecules as compared to other monocyte subsets (**Fig. 5D**). We also found higher frequencies of activated memory T cells expressing high levels activation markers (**Fig. 5F-G**), which future studies should investigate for their potential to recognize microbiota-derived antigens or provide enhanced immunosurveillance against pathogens. Another immune population associated with IMCV was CD69^high^ activated MAIT cells, consistent with reports of their activation by microbiome-derived molecules.^57^ Recent studies have found that MAIT cells can confer protection against colitis and inflammation-induced colorectal cancer after activation by microbiome-derived ligands.^61,62^

The most prominent immune variation associated with microbiome features in our study was the tonic-IFN response. Since many ISGs encode antiviral mediators, our findings suggest that the microbiome could be a source of interindividual variability in susceptibility to infection. This is consistent with studies that have shown that reduced microbiome diversity or depletion by antibiotics result in poor antiviral protection and weakened immune responses.^19^ Additionally, the IFN axis plays a critical role in immunomodulatory therapies, which often exhibit substantial variability in patient response.^12^ We observed an elevated tonic-IFN state in individuals with stronger immune responses to the *influenza* vaccine but also in patients treated with anti-PD1 checkpoint blockade who only responded upon IFN-axis inhibition.

While beneficial and detrimental microbial communities have been identified in animal models, specific findings have not always applied to humans. Additionally, widespread discrepancies between studies make the taxonomic definition of a “good” human microbiome elusive.^63^ Instead, there is growing recognition that microbiome genetic content that encodes functional potential may matter more than the phylogenetic identity of the bacteria. Our findings identify a combination of microbial pathways and molecules that associate with tonic antiviral activity. These include SCFAs, polyamines, microbiome-derived LPS endotoxin and peptidoglycans, and primary bile acids, consistent with mechanistic studies in model organisms showing that commensal bacteria contribute to shaping the immune system.^19,64–68,40,51,52,69–71^ Notably, many of these immunomodulatory molecules were elevated in the same subset of individuals, suggesting that their production may be coordinated with one another and that their combination may matter. Future studies should focus on revealing the causal relationship between the variable features identified in this study and their impact on disease development or protection.

We envision that the data resource presented here will provide insights and context for ongoing and future efforts to modulate the microbiome and the immune system for therapeutic benefit through various approaches such as dietary interventions,^72–74^ microbial colonization,^75^ or microbiome-informed precision immunotherapy.

## Limitations

The ImmunoMicrobiome study profiled biological variation in a cohort of study participants living in the San Francisco Bay Area at the time of the study. While the data does not account for the biological variability exhibited across the whole human species, the local sampling did enable us to exclude large environmental variables associated with geography as main drivers of the inter-individual variation we describe. Similarly, ImmunoMicrobiome specimens were collected within a 3-month period, which both limited the impact of known seasonal influences as well as our ability to examine these effects on the systems studied. Larger cohorts and longitudinal sampling will be required to address the impacts of geography and seasonality that were not examined in this study.

## Methods

### Study participant cohort: Subject enrollment and consent

Healthy subjects were enrolled and written informed consent was obtained under the UCSF IRB approved protocol #18-27022. Healthy status and eligibility were defined through screening questionnaires, based on the absence of cancer history, autoimmune disease, severe allergy or asthma, known immunodeficiency or chronic infection. Study participants were major, non-pregnant and free of medical treatment affecting the immune system or the microbiome at the time of enrolment and during the study. Specifically, history of surgery, use of immunosuppressants or broad spectrum antibiotics taken during or in the last 4 months before enrolment were an exclusion criteria, and treatment history has been documented. Study participants who underwent seasonal vaccines before the study visit were at least 30 days out of their last shot at the time of sampling and vaccine history has been accounted for in the analysis. Most study participants lived in San Francisco County (except for few who resided in Alameda and San Mateo counties). The race and ethnicity bias in the cohort, characterized by overrepresentation of non-Hispanic White and Asian individuals, reflected the demographic breakdown of the counties where participants lived (source: http://data.census.gov).

### Biospecimens collection and processing

Stool samples by study participants at the research facility within a 6h window for all participants. The samples were transferred to the research team and put on ice until processing. The samples were manually homogenized and aliquoted under a biosafety cabinet, and cryopreserved at -80 degree Celius.

Blood was collected in EDTA tubes within a 90 min window for all participants of the study. Plasma samples were obtained by centrifugation of whole blood samples at 2000g for 15 minutes at 4°C and aliquots were cryopreserved at -80 degree celsius. Peripheral blood mononuclear cells (PBMC) were obtained by density gradient centrifugation of whole blood at 800g for 30 min at room temperature without brake, using Ficoll-Paque (Cytiva). Isolated cells were either further processed for mass cytometry preparation or cryopreserved in a solution containing fetal Bovine Serum and 10% DMSO (Sigma) for further processing in CITE-Seq preparation.

### Survey data collection and preprocessing

Survey data was collected through self-administered questionnaires. Short term dietary data was collected through the NIH validated Automated Self-Administered 24-Hour (ASA24®) Dietary Assessment Tool (https://epi.grants.cancer.gov/asa24). General health, diet, medical history and demographic data was collected through a questionnaire designed by the research team.

The nutrient levels from ASA24 survey were collected, features with zero values in all samples were removed, and the data was log-transformed and scaled for the integrative analysis.

### Microbiome profiling

#### Whole metagenome shotgun sequencing DNA extraction

Frozen stool samples were thawed for bacteria lysis and DNA extraction performed using a combination of the cetyltrimethylammonium bromide (CTAB) chemical, phenol/chloroform solvents and bead beating mechanical forces. Briefly, 200 mg of human stool samples were incubated at 65 degree celsius for 15 minutes in a 5% CTAB solution. A phenol/chloroform solution was added and the samples were homogenized with bead beating for 2 rounds of 30 seconds in a Fastprep24 homogenizer (MP Biomedicals) using Lysing Matrix E beads (MP Biomedicals). The aqueous phase was collected after centrifugation and mixed with chloroform before centrifugation. Supernatant was then mixed with a 30% PEG solution and incubated overnight at 4°C. Precipitated DNA was collected after 2 rounds of washing in ice cold 70% Ethanol. Magnetic bead-based DNA cleanup was then performed using AMPure XP beads (Beckman Coulter).

#### Whole metagenome shotgun library construction and sequencing

Whole metagenome shotgun sequencing libraries were generated using the Nextera XT (Illumina). Custom 12-bp dual unique indices were introduced during PCR amplification. Libraries were pooled at the desired relative molar ratios and DNA cleanup was performed using AMPure XP beads (Beckman Coulter) for buffer removal and library size selection. Sequencing reads were generated using a NovaSeq S4 flow cell.

#### Whole metagenome shotgun data preprocessing

Sequencing reads were processed with a custom pipeline. Briefly, reads were quality-filtered and adapters were removed using FastP v0.23.2^76^ (ref https://doi.org/10.1002Imt2.107) with standard parameters. Human reads were filtered out of the dataset using the computational tool BMTagger (ftp://ftp.ncbi.nlm.nih.gov/pub/agarwala/bmtagger/). BioBakery tools Metaphlan version 3.1 was used to estimate taxa community and HUMAnN3^77^ was used to estimate gene family/pathway abundances (units of length-normalized average copy numbers of genes summarized at the gene family or pathway level). For MOFA analysis, the species abundance data from MetaPhlan3 was normalized by the total abundance per sample (after removing ‘UNKNOWN’ category), the species with non-zero values in more than 10 samples were retained, and the data was centered log-ratio (CLR) transformed and scaled for the integrative analysis. Gene family and pathway abundances were normalized to average copy numbers (reads per kilobase per genome equivalent, RPKG) using MicrobeCensus v1.1.1 (ref http://genomebiology.com/2015/16/1/51). For MOFA analysis, the pathway abundance data from HUMAnN3, the features with non-zero values in more than 10 samples were retained, and the data was log-transformed and scaled for the integrative analysis. For PCA analysis, when unnormalized microbiome composition data was used to examine factors’ association with the microbiome, abundances of the most prevalent species across the cohort were included as an input to mitigate the effect of species sparsity on PCA.

#### Species level contribution to gene family and pathway abundances

Species-level contributions to gene family abundances were calculated on the species-stratified output of HUMAnN3. The prevalence represents the percentage of the cohort that carried a non-zero copy number for a given gene family in a given species. Thus, species “contribution to a gene family” is the median across the cohort of the ratio of the copy number of genes for a given species over the total copy number of genes for all species (RPKG fraction). The same metrics were calculated for Species-level contributions to pathway abundances, using gene family copy numbers summarized at the pathways level.

### Metabolome profiling

#### Targeted and untargeted metabolomics sample preparation

Frozen stool samples were later lyophilized over 48 hours, crushed into a fine powder, 25 milligrams of powder was weighed out per study participant into Precellys tissue homogenizing tube, and samples were stored back at -80°C. Frozen plasma samples were thawed on ice, centrifuged at 15,000g for 5 minutes at 4°C. An equal volume of cell-free supernatant was taken from each study participant for extraction. Metabolites were extracted from each biospecimen by addition of 4 volumes of extraction solvent (1:1 acetonitrile and methanol with 5% LCMS grade water). Mixture was vortexed for 1 minute then incubated at -20°C for 30 minutes. Upon completion of mixing and incubation, samples were centrifuged at 15,000g for 30 minutes at 4°C. Supernatants were extracted and stored at -20°C until time of analysis.

#### Very short chain fatty acids targeted metabolomics

Samples were extracted using a biphasic extraction using 0.5 mL H20 with Internal standards, 0.1mL concentrated HCL and 1 mL MTBE. The samples are shaken for 30 min at room temperature then centrifuged for 2 min at 14000 rcf. 100 uL of the organic phase is aliquoted into a glass crimp vial with 25 uL of MTBSTFA. Vial is crimp caped and shaken at 80°C for 30 min. After derivatization vial is placed in GC sampler for analysis. 1 uL of derivatized sample is injected using a split method at an inlet temperature of 250°C. A constant flow of 1.2 mL/min using Helium is used throughout the run. The oven temperature program goes as follows starting at 50°C for 0.5 min then ramping to 70C at 5°C/min holding for 3.5 min, ramp to 120°C at a rate of 10°C/min, finally ramping to 290°C at a rate of 35°C/min holding for 3 min for a final run time of 20.857 minutes. The transfer line is set to 290C while the EI ion source is set to 250C. The Mass spec parameters collect data from 85m/z to 5m/z at an acquisition rate of 17 spectra/sec. A 6-point Curve with internal standards is injected alongside the samples and used for absolute concentration as well as a pool sample used as QC which is injected every ten samples. Mass hunter Quant is used for data processing and picking of target peaks. Then normalized to the amount of sample injected.

#### Primary Metabolism untargeted metabolomics

Samples extracted using 1mL of 3:3:2 ACN:IPA:H2O (v/v/v). Half of the sample was dried to completeness and then derivatized using 10 uL of 40 mg/mL of Methoxyamine in pyridine. They are shaken at 30C for 1.5 hours. Then 91 uL of MSTFA + FAMEs to each sample and they are shaken at 37C for 0.5 hours to finish derivatization. Samples are then vialed, capped, and injected onto the instrument. We use a 7890A GC coupled with a LECO TOF. 0.5 uL of derivatized sample is injected using a splitless method onto a RESTEK RTX-5SIL MS column with an Intergra-Guard at 275C with a helium flow of 1 mL/min. The GC oven is set to hold at 50C for 1 min then ramp to 20C/min to 330C and then hold for 5 min. The transferline is set to 280C while the EI ion source is set to 250C. The Mass spec parameters collect data from 85m/z to 500m/z at an acquisition rate of 17 spectra/sec.

#### Biogenic amines untargeted metabolomics

(to be completed by Oliver’s Team)

#### Metabolomics data preprocessing

The data from targeted metabolomics for very short chain fatty acids, and untargeted metabolomics for biogenic amines and primary metabolites were concatenated. The features with unknown annotation were discarded and duplicate features based on the same name or same InChiKey were deduplicated by selecting the feature with lowest variation across the batch-control samples. The data was log-transformed and pareto scaled (scaled data multiplied by square-root of standard deviation) for the integrative analysis.

### Immune profiling CyTOF

#### Mass-tag antibody conjugation and antibody cocktail preparation for mass cytometry

Mass cytometry antibodies were prepared using the MaxPAR antibody conjugation kit (Standard Bio Tools) according to the manufacturer’s instructions. Following conjugation, antibodies were diluted at concentrations ranging from 0.2 and 6 mg/mL in Candor PBS Antibody Stabilization solution (Candor Bioscience GmbH, Wangen, Germany) supplemented with 0.02% NaN3 and stored at 4°C. Each antibody was titrated to identify optimal staining concentrations using human PBMCs. A surface staining antibody cocktail and an intracellular antibody cocktail made of all the 2 types of target were prepared for the staining of all batches of the study. A summary of all mass cytometry antibodies, metal tags and concentrations used for analysis can be found in the STAR methods.

#### Mass cytometry sample preparation

Briefly, PBMCs were incubated immediately after isolation with a viability staining solution containing 50uM of cisplatin (Sigma-Aldrich) for 1 min at room temperature and fixed in a solution containing 1.6% Paraformaldehyde (PFA) (Electron Microscopy Science) and cryopreserved in a solution containing 0.5% BSA and 10% dimethyl sulfoxide (DMSO). Fixed cells were later thawed and incubated with benzonase nuclease (Sigma-Aldrich), permeabilized with a solution containing 0.2% saponin, barcoded by incubation with a combination of custom palladium metal stable isotopes for 15 minutes at room temperature, and pulled in batches of 20 samples. Fc receptors were blocked using Human TruStain FcX (Biolegend) and cells were stained with custom metal ion-labeled antibody cocktails. Extracellular staining was performed at room temperature for 30 minutes on a shaker set on 90 RPM. Intracellular staining was then performed using eBioscience Permeabilization Buffer (Invitrogen) at 4°C for 60 minutes on a shaker set on 90 RPM. Cells were later incubated with Cell-ID Intercalator-Ir (Standard Biotools) at 0.0625 uM and 4% PFA overnight at 4°C. Before running, cells were resuspended in cell acquisition solution (Standard Biotools) that contained EQ Four Element Calibration Beads (Standard Biotools). Except for permeabilization steps and final steps, washes were performed using a cell staining solution containing PBS,BSA and EDTA or PBS.

#### Mass cytometry data acquisition

Mass cytometry sample pools were diluted in Cell Acquisition Solution containing bead standards to approximately 1 x 10^6 cells/mL and then analyzed on a Helios mass cytometer (Standard Biotools) equilibrated with Cell Acquisition Solution. A minimum of 10 x 10^6 cell events were collected for each barcoded set of samples at an event rate of 400 events/second.

#### Mass cytometry data normalization and de-barcoding

Bead standard data normalization and de-barcoding of the pooled samples into their respective conditions was performed using the R package from the PICI institute available at https://github.com/ParkerICI/premessa.

#### Mass cytometry data batch correction, clustering, annotation, visualization, and differential protein expression analysis

The mass cytometry data for events corresponding to live cells were extracted from bead-normalized FCS files, arcsinh-transformed using a cofactor of 5, and subjected to batch-correction using cyCombine ^78^ (v0.1.5). The dimensionality reduction and unsupervised clustering analysis were performed to obtain 225 clusters of single cells using cyclone ^79^, which uses FlowSOM ^80^ (v2.0.0) for clustering, and uwot R package (v 0.1.10) for dimensionality reduction. The clusters were annotated manually using known marker expression patterns, and were visualized using SCAFFoLD ^29^ (v0.1; default parameters ). For the landmark populations used as input to SCAFFoLD, an equal number of events from all batch-corrected FCS files were combined to produce a concatenated dataset of 100k cells, which was manually gated on CellEngine to produce FCS files of populations of known phenotype. The SCAFFoLD map validated the marker-based manual annotation of individual clusters. The cluster frequencies were normalized by the total number of cells per sample, log-transformed, and scaled for the integrative analysis.

The differential protein expression analysis was performed using Kruskal-Wallis test to identify markers that are different between groups of CyTOF clusters. The clusters were grouped into two to three groups based on positive or negative association or no association with a MOFA factor (adj. p < 0.1 from linear models - see below). Through 10 iterations, 100 cells were randomly drawn from each of the cluster groups, and the normalized protein expression was compared between the groups using Kruskal-Wallis test. The markers that were significantly different (p < 0.00001) in all 10 iterations were identified as significantly differentially expressed proteins and are presented in Fig. 4D, 4H, and S4D-G). Pairwise comparisons were performed as a post-hoc test to identify pairs of cluster groups with differential expression and a significance score was calculated based on (-log10)p-values, scaled to the highest value.

### Immune profiling CITE-seq

#### Processing of PBMCs for cellular indexing of transcriptomes and epitopes by sequencing (CITE-seq)

Cryopreserved PBMCs were thawed. Cell counts and viability was determined using the CellacaMX. PBMCs were pooled in batches of sixteen samples in equal numbers to a total of 1 million cells. Pools were resuspended in Cell Staining Buffer (Biolegend) and incubated with Fc Receptor Blocking Solution (Biolegend) at 20mg/mL for 10 minutes on ice. Cells were then stained with a cocktail of 140 TotalSeq-C oligo-tagged antibodies (Biolegend) for 30 minutes at 4°C. Stained cells were washed in PBS containing 1% BSA, filtered through a 70uM filter, and re-suspended in a final solution of PBS containing 0.04% BSA. Cells (n=60,000) were loaded on the Chromium Controller (10X Genomics) for GEM generation. The cDNA libraries were generated using the Chromium Next GEM Single Cell 5’ v1.1 Library Kit for gene expression (GEX) and the Chromium Single Cell 5’ Feature Barcode Library kit for antibody derived tag (ADT) according to manufacturer’s instructions (10X Genomics). Each batch of sixteen samples was sampled four times to prepare four independent cDNA libraries per batch to obtain a sufficient number of cells per sample. The libraries were subsequently sequenced on a NovaSeq 6000 S4 platform (Illumina).

#### Bulk RNA library preparation and data analysis for demultiplexing of CITE-seq libraries

Cryopreserved PBMCs were thawed. Cell counts and viability were determined using the CellacaMX. 0.5 million cells were used for each sample. RNA was extracted using the Quick-RNA MagBead kit (Zymo Research) on the KingFisher automated extraction and purification system (ThermoFisher) according to manufacturer’s instructions. Quality and quantification of extracted RNA was assessed on the 5300 Fragment Analyzer System (Agilent). Next, cDNA libraries were generated using the Universal Plus mRNA-Seq with NuQuant kit according to manufacturer’s instructions (Tecan). The libraries were subsequently sequenced on a HiSeq 4000 platform (Illumina).

Sequencing reads were aligned to the human reference genome and Ensembl annotation (GRCh38 genome build, version 95) using STAR (v2.7.5c) ^81^with the following parameters: --outFilterType BySJout - -outFilterMismatchNoverLmax 0.04 -outFilterMismatchNmax 999 --alignSJDBoverhangMin 1 -- outFilterMultimapNmax 1 -alignIntronMin 20 --alignIntronMax 1000000 --alignMatesGapMax 1000000. Duplicate reads were removed using Picard Tools (v2.23.3) (http://broadinstitute.github.io/picard/). Nucleotide variants were identified from the resulting bam files using the Genome Analysis Tool Kit (v4.0.11.0) following the best practices for RNA-seq variant calling ^82,83^. This included splitting spliced reads, calling variants with HaplotypeCaller (added parameters: --dont-use-soft-clipped-bases -standcall-conf 20.0), and filtering variants with VariantFiltration (added parameters: -window 35 cluster 3 --filter-name FS -filter FS > 30.0 --filter-name QD -filter QD < 2.0). Variants were further filtered to include a list of high quality SNP for identification of the subject of origin of individual cells by removing all novel variants, maintaining only biallelic variants with MAF greater than 5%, a max missing of one individual with a missing variant call at a specific site and requiring a minimum depth of two (parameters: --max-missing 1.0 --min-alleles 2 --max-alleles 2 --removeindels --snps snp.list.txt --min-meanDP 2 --maf 0.05 -- recode --recode-INFO-all –out).

#### Sequence alignment and generating raw count matrix

The raw sequencing reads from gene expression and surface protein expression libraries were aligned to human genome reference (vGRCh38-2020-A) with annotations from Gencode (v32-primary assembly) and to the TotalSeq-C antibody reference panel (BioRad), respectively, using cellranger (v6.0.2) with default options.

#### CITE-seq data demultiplexing and quality control

The raw counts of gene expression from cellranger output were imported and analyzed using Seurat^84^ (v4.0.3). Because each library of 10X data contained cells from sixteen samples, we first performed demultiplexing of the data based on SNP profiles to map individual cells to its sample of origin. Specifically, freemuxlet (vAug2021; doi:10.1101/2023.05.29.542756) was used to cluster the cell barcodes based on concordant SNP profiles (list of high-quality reference SNPs obtained from the 1000 ^85^Genomes Consortium; ), and the SNP profiles of freemuxlet clusters were mapped to the genotypes of study participants using bcftools ^86^(v1.10.2), which was generated from the bulk RNA-seq data, allowing us to map each cell barcode to the sample of origin. The *inter-sample* doublets (DBL: barcodes containing mixture of multiple SNP profiles) and empty droplets (AMB: barcodes lacking sufficient coverage of reference SNPs for freemuxlet clustering) were removed, and the barcodes with SNG calls (droplets containing a single cell or multiple cells from a same individual (intra-sample doublets)) were subjected to further filtering. DoubletFinder was used^87^ (v2.0^87^) to filter barcodes containing heterotypic intra-sample doublets (i.e. the droplets containing multiple cells of different cell-types from a same participant) as described here https://github.com/chris-mcginnis-ucsf/DoubletFinder. Briefly, first the data was transformed using SCTransform, principal component analysis (PCA) was performed using npcs=35, nearest neighbors were computed using FindNeighbors(), and the data was clustered using FindClusters() in Seurat. The PC neighborhood size (pK) was adjusted for each library separately by performing a pK-pN parameter sweep using paramSweep_v3() of DoubletFinder package. The proportion of expected homotypic doublets were estimated based on cluster assignments from FindClusters() output, and the number of expected heterotypic doublets were determined using the inter-sample doublets (DBL calls from freemuxlet) by dividing the fraction of inter-sample doublets by the number of all possible combinations of inter-sample doublets, otherwise the default options were used. The barcodes annotated as heterotypic doublets using DoubletFinder were discarded and the remaining barcodes representing single-cells were subjected to further quality control. The barcodes containing less than 100 genes and genes that are detected in less than 3 cells were filtered out. We further filtered out the barcodes based on library-specific cutoffs for the number of genes per cell (nFeature_RNA < 300-800 & > 6,000), the number of unique molecular indexes (UMIs) per cell (nCount_RNA < 400-2,000 & > 40,000), mitochondrial content (>15%), and ribosomal content (<8-15% & >60%) to remove poor quality cells. The resulting high-quality cell-barcodes were used for further analysis.

#### Normalization, dimensionality reduction, clustering, and cluster annotation

The gene expression data from all libraries were merged, log-normalized, and scaled with regressing out for RNA counts per cell, mitochondrial and ribosomal contents, and cell-cycle states. Top 2,000 most variable features were found using the ‘vst’ method, PCA was performed using npcs=50, and batch-effect (library-specific bias) was normalized using Harmony ^88^ (v1.0;).

The ADT data was normalized and denoised using ‘denoised and scaled by background’ (dsb) method (^89^) for each library separately. The dsb utilizes protein-specific background signal from empty droplets and signal from isotype control antibodies to accurately estimate and correct for cell-to-cell technical noise in each cell. The ADT data was normalized and denoised using cell barcodes containing less than 100 RNA counts and more than 10 protein counts as empty droplets and expression of five isotype antibodies included in the TotalSeq-C panel as isotype-control signal in DSBNormalizeProtein() from dsb package^89^ (v0.3.0). The normalized values smaller than -10 were truncated by substituting them with zero. The expression data for the isotype control antibodies were excluded from the further analysis. The protein expression data demonstrated the largest batch-effect; therefore, we used reciprocal PCA (RPCA) to integrate the protein data. Briefly, after scaling the data and calculating PCs for each library, integration anchors were calculated using the first library from each batch as reference, and then the data was integrated using IntegrateData() function in Seurat. The integrated protein data was scaled, the ADT counts per cell and cell-cycle states were regressed out, and PCA was performed using npcs=30.

Weighted nearest neighbor (WNN) graph was constructed using first 30 harmony dimensions for gene expression, first 18 integrated PCs for protein expression, and prune.SNN = 1/20. Using the WNN graph, dimensionality reduction was performed using Uniform Manifold Approximation and Projection (UMAP) and 56 clusters of cells were identified using smart local moving (SLM) algorithm and resolution=2 using Seurat’s FindClusters function. Marker genes and proteins were identified for each cluster using Model-based Analysis of Single Cell Transcriptomics (MAST) algorithm (https://github.com/RGLab/MAST/) with library as a latent variable. The clusters were assigned cell type annotation using the expression of canonical marker genes and proteins as depicted in Fig. S1A. A representative sample of the data was used for UMAP coordinate visualizations in Fig. 1F and S1B.

Gene expression variance in the ImmunoMicrobiome cohort, the single-cell data for cell types were isolated, and subjected to PCA, and batch-correction using harmony. The top ten dimensions, capturing the major axes of variation, were selected, the dimension coordinates of single-cells were averaged per participant, and these averages were depicted in Fig. 1G for monocytes. Linear models were used to identify the genes associated with averaged Dimension 1 coordinates.

#### Preparing data for integrative analysis

For each cell type, we used pseudo-bulked gene expression per sample in the integrative analysis. The cell types that contained more than 20 cells per sample in more than 10 samples were kept for further analysis. The normalized single cell gene expression was averaged(in linear space) across samples to generate a sample-by-gene matrix for each cell type. To remove the lowly expressed genes, the genes that were detected in more than 30% of samples were retained. The pseudo-bulked expression data was batch-corrected using ComBat from sva package^90^ (v3.40.0), the residual batch-effects were eliminated by removing genes associated with batches after ComBat (Kruskal-Wallis adjusted p-value < 0.01), and the depletion of batch-effects were validated using principal variance component analysis (PVCA). The top 5,000 most variable genes identified using median absolute deviation (MAD) were selected, and the expression data for these genes were log-transformed and scaled for the integrative analysis.

The cell type frequencies were normalized by the total number of cells per sample, log-transformed, and scaled for the integrative analysis.

The ADT data was also filtered in the same manner as gene expression data and was batch-corrected using ComBat. This data was used for follow up analysis of MOFA factors.

#### Enrichment analysis

Enrichment analysis was performed using minimum hypergeometric tests (mHG)^91^ to identify hallmark signatures^92^ that are significantly enriched (BH-adjusted p < 0.1) in the significantly associated features (BH-adjusted p < 0.1 and |feature weight| > 0.1; see Evaluating composition of latent factors) for each factor in each dataset. The features were ranked based on factor weights for the enrichment analysis. The selection of an appropriate background set of features is crucial for the accurate enrichment analysis, therefore the features that were detected in our assays were used as background in the mHG test. To calculate factor-specific signature scores, the log-transformed and scaled expression of core genes (genes driving the mHG enrichment) were weighted by MOFA weights and summed for each participant (used in Fig. 4A). The importance of signatures within each factor was evaluated by combining enrichment p-values of each signature across all datasets using the Cauchy combination test, which is known to be robust to the dependence structure underlying p-values to be combined ^93^. The joint p-values were BH-adjusted to obtain joint adj.p. Several hallmark signatures were significantly enriched in most factors. We noticed that the first five factors were associated with similar immune signatures. To select the hallmark signatures for the follow up investigation, we took the top 5 most significant signatures (based on the joint adj.p) for each of the first five factors and counted the number of CITE-Seq cell subsets they were significantly enriched in (Fig. S2D). Based on this analysis, we selected these hallmark signatures to characterize the factors: Inflammation-related gene sets - inflammation response (M5932), TNF alpha signaling via NFKB (M5890) and TGF beta signaling (M5896); Interferon-related gene sets - IFN alpha response (M5911) and IFN gamma response (M5913). Given large overlap in features between these hallmark signatures, beyond the initial analysis in Fig. 1-2 overlapping genes between IFN-related and inflammation-related signatures were excluded from the gene sets (Fig. 3-7). Additionally, gene sets related to tonic-IFN ^27^ and, prolonged-IFN ^33^ responses were derived from previous publications.

For comparison between high-IMCV and high-IIV sub-cohorts, limma ^94^was used to identify genes significantly differentially expressed between the two groups (BH-adjusted p < 0.1). Enrichment analysis was performed using mHG to identify the gene signatures used in the study that are significantly enriched (BH-adjusted p < 0.1) in high-IMCV or high-IIV sub-cohorts in each dataset. The features were ranked based on log2 fold-difference between high-IMCV and high-IIV sub-cohorts for the enrichment analysis. To calculate the composite signature scores for each individual, the log-transformed and scaled expression of the core genes driving the enrichment of a given signature were averaged using median for each individual.

#### Serological profiling

Serological profiling data was generated by Proximity Extension Assay technology using oligonucleotide labeled antibody with the Target 96 Inflammation Panels.

### Integrative analysis

#### Building MOFA model

As described above, all omics datasets were standardized by log-transformation and scaling (by calculating Z-scores). The Multi-omics Factor Analysis (MOFA) ^31^ (v1.2.2) was used to extract the shared variation across a total of 42 datasets (an ASA24 data, a metabolomics dataset, two microbiome datasets, two immune frequency datasets, and 36 gene expression datasets). MOFA performs matrix decomposition to represent the multi-omics data in reduced dimensions as represented by latent factors that capture biological and technical sources of variation. Our evaluation of MOFA models with different factor numbers revealed that the 15-factor model captured a substantial amount of shared variation between immune and microbiome compartments.

#### Evaluating composition of latent factors

To evaluate the composition of factors, the multi-omics features significantly associated with factors were identified using linear regression models (BH-adjusted p < 0.1 and |feature weight| > 0.1), while controlling for experimental batches, if available.

#### Quantification and Statistical Analysis

Most statistical analyses were carried out in R. The p-values were adjusted for multiple-testing using Benjamini-Hochberg procedure, unless mentioned otherwise. The correlation analysis was performed using Spearman’s rank method unless indicated otherwise. Significance of differences in pairwise comparisons was performed using Wilcoxon rank sum test and in multi-group comparisons using Kruskal-Wallis test. Linear regression models were used to find features associated with MOFA factors.

Correlations were visualized by a line using the method lm in ggplot2 geom_smooth but correlations were calculated using the package cor.test.

## Acknowledgments

This study was supported by the Bakar ImmunoX Initiative, the Benioff Center for Microbiome Medicine, National Microbiome Initiative, American Association for Cancer Research award 20-20-01-SPIT, Cancer Research Institute award CRI4437, NIH awards R01DE032033, DP5OD023056, R01HL122593 (P.J.T.), R01CA255116 (P.J.T.), and R01DK114034 (P.J.T.), American Cancer Society award RSG-22-141-01-IBCD, DOD US Army Med. Res. Acq. Activity Award BC220499 to M.H.S. The CyTOF instrument used in this study was purchased with assistance by NIH award S10 OD018040. C.N. was supported by the National Institutes of Health (F32GM140808). P.J.T is a Chan Zuckerberg Biohub-San Francisco Investigator.

We thank the Chan Zuckerberg Biohub Microbiome Initiative for library constructions and sequencing, Sue Lynch, Moriah Sandy, Kyle Spitler and The Benioff Center for Microbiome Medicine for providing protocols, guidance and equipment for stool sample processing and metabolomics. Andy Minn and Divij Matthew for providing the cancer immunotherapy validation dataset. Les Dethlefsen and Lloyd Bod for guidance and advice, Daryll Gempis & Elizabeth Joyce from the UCSF microbiology department for facilitating study participant sampling collection visits. Tara Taeed and Laura Trupin from UCSF Rheumatology department for phlebotomy sample collection support, the CoLabs, the Bakar ImmunoX Initiative, UCSF CAT Core for sequencing and UCSF Flow Core and Stanley Tamaki for CyTOF instrumentation support. UCSF CAT is supported by UCSF PBBR, RRP IMIA, and NIH 1S10OD028511-01. UCSF Parnassus Flow CoLab is supported by RRID:SCR_018206 and DRC Center Grant NIH P30 DK063720. We thank all the study participants of the ImmunoMicrobiome cohort.

## Author contributions

Clinical study design: J.B. and M.H.S.; Specimen collection and processing: J.B., B.D., M.R, I.T., D.M., A.S., M.C., V.J., C.E.; Data generation: J.B., B.D., B.Y., O.F.; Conceptualization: J.B., R.K.P., G.F., M.H.S.; Data Curation: J.B., R.K.P.; Formal Analysis: J.B., R.K.P, B.D.,C.N., J.E.B., P.J.T.; Funding Acquisition: J.B., G.F.,O.F., M.H.S., P.J.T.; Investigation: J.B., R.K.P.; Methodology: J.B., R.K.P.,C.N., A.C., G.F., M.H.S., P.J.T.; Project Administration: J.B., A.C., G.F., M.H.S.; Resources: J.B., A.C.,B.Y., O.F, P.T.,E.E.M., G.F., M.H.S.; Software: J.B., R.K.P., C. N., J.E.B.; Supervision: J.B., A.C., G.F., M.H.S., P.J.T.; Validation: J.B., R.K.P., C.N., G.F., M.H.S.; Visualization: J.B., R.K.P.; Writing – Original Draft Preparation: J.B.; Writing – Review & Editing: J.B., B.D., R.K.P., C.N., J.E.B., G.F., M.H.S., P.J.T.

## Supplemental methods

### CyTOF annotation

Immune populations were annotated based on their expression of the following marker. B cells were CD19+ and CD27 was used to identify naive and memory subsets. T cells were CD3+. Within the T cells, CD56 was used to identify NKT cells and gdTCR was used to distinguish gdT cells and abT cells. Within abT cells, CD4 and CD8 were used to identify lineages, CCR7 and CD45RA were used to identify naive and memory cell subsets. NK cells were CD3-HLA-DR low/- and CD56+. NK subsets were defined using CD16. Myeloid cells were CD3-CD19-CD56-CD123-HLA-DR+. Within the myeloid, monocyte subsets were identified by CD14 and CD16, conventional dendritic cells were CD14- and CD16- and subsets were defined using BDCA3 CADM1 Clec9A BDCA1 SIRPa and Axl. pDCs were HLA-DR+ and CD123+ and Axl was used to define subsets. Residuals neutrophils were identified as CD15+ CD16+ and residuals basophils as HLA- DRlow/- and FcER1A+ and CD123 int.

---

**Figure S1.**
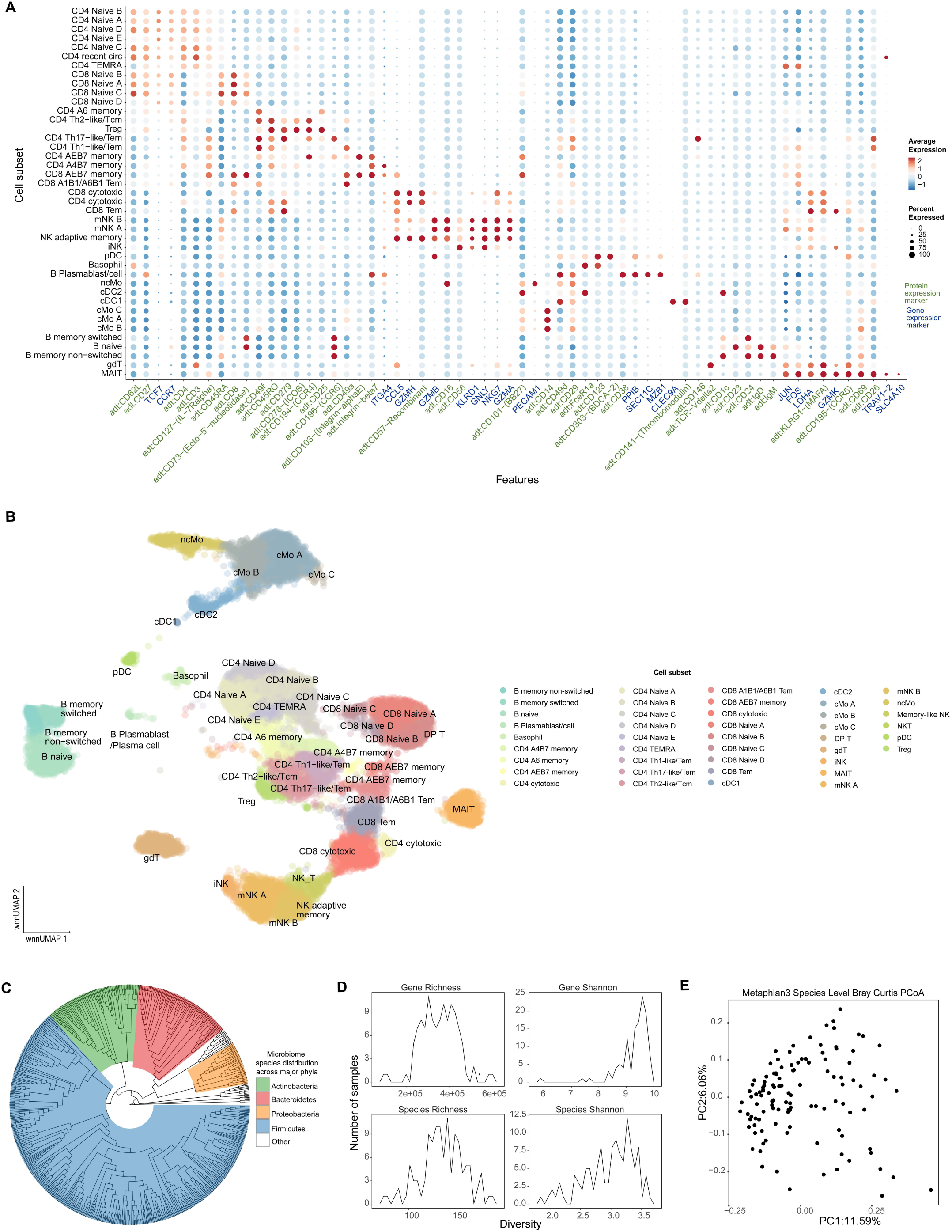
Related to Figure 1. A. Dot plot quantifying the prevalence and expression of key marker genes and surface proteins used to annotate CITE-Seq clusters. Marker names are colored based on the library source. Gene expression library markers are annotated in blue. Surface protein expression markers from the antibody derived tag library are indicated in green. B. UMAP visualization of CITE-Seq data. Single cell mapping and coloring is the same as Fig. 1F. Annotation details unaggregated cell subsets identified by unsupervised clustering. C. Phylogenetic tree of species identified in the ImmunoMicrobiome cohort. Colors represent major microbiome phyla. D. Line plots describing microbiome diversity across the cohort. E. Scatter plot showing the distribution of the study participants based on their microbiome composition at the species level after dimensionality reduction using PCoA with Bray-Curtis dissimilarity distance metric.

**Figure S2.**
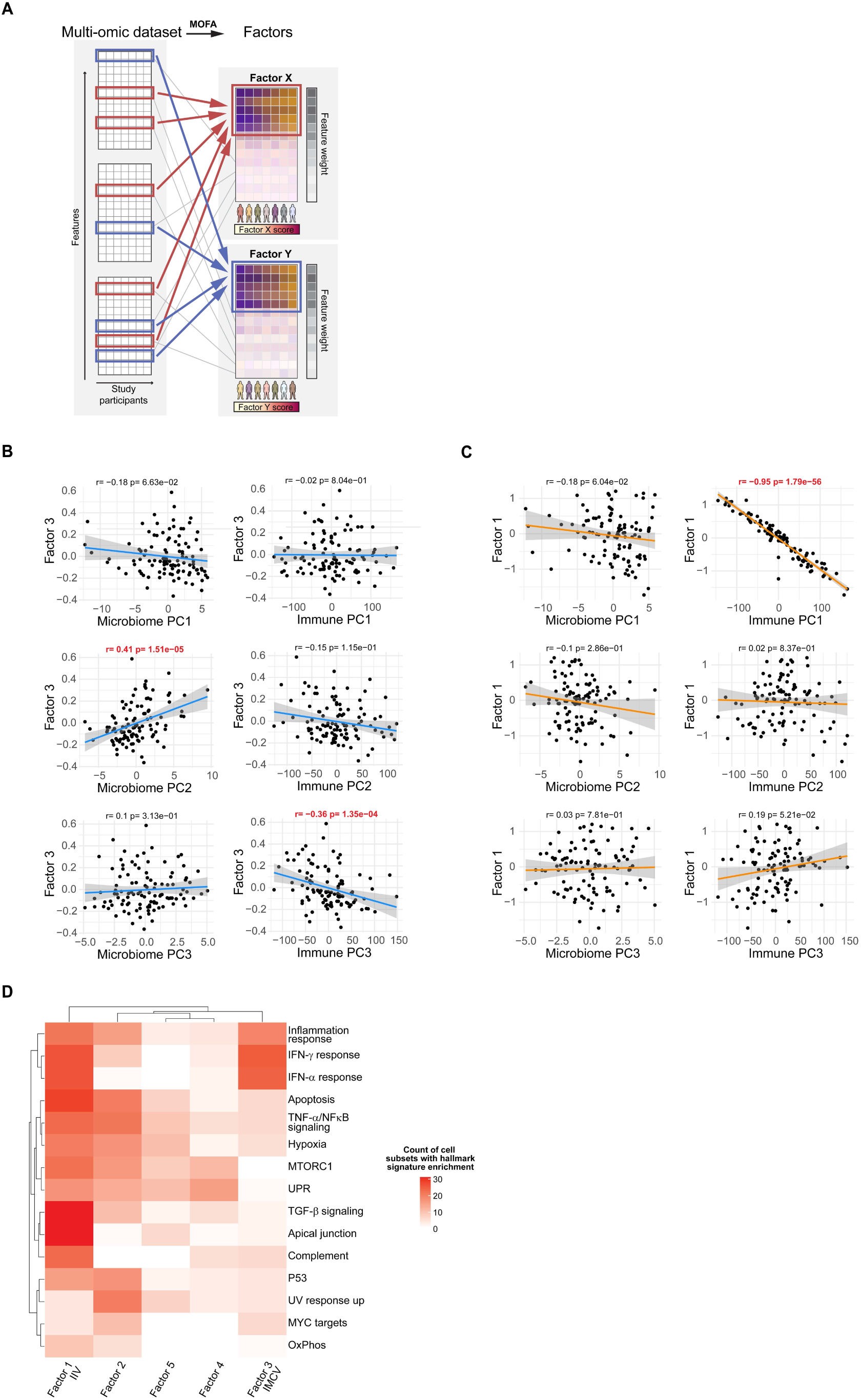
Related to Figure 2. A. Schematic of integrative approach. Features that vary concomitantly across datasets in the cohort are captured by factors. For each factor, features are ordered based on their weight on the factor (feature weight) and study participants are ordered based on factor values (factor score). In the schematic, features driving the variation captured by factors are indicated with red and blue boxes. B-C. Scatter plots of factor scores and first three principal components from the microbiome species relative abundance data (Microbiome PC1-3) and CITE-Seq gene expression data (Immune PC1-3). Factor 3 is shown in panel B and Factor 1 in panel C. D. Heatmap quantifying the number of CITE-seq cell subsets that have significant enrichment of signatures among the first five MOFA factors. Signatures were evaluated using mHG test and hallmark gene set library.

**Figure S3.**
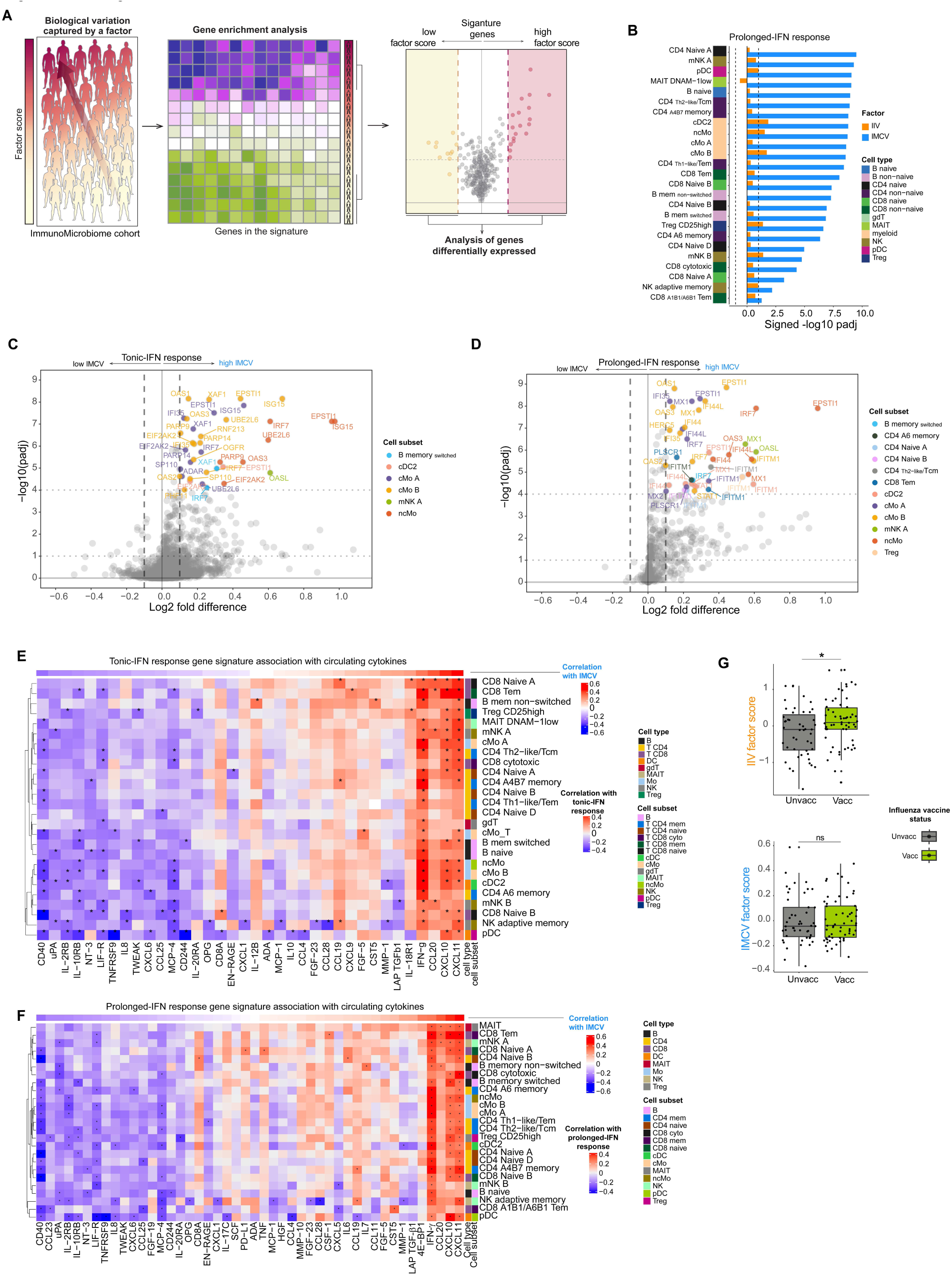
Related to Figure 3. A. Schematic of the analytical approach used in figure 5 and 6. Enrichment analysis is performed within MOFA factors, to assess association between factor scores and transcriptional signatures of interest. For signatures enriched, differential gene expression comparing study participants with high factor scores to those with low factor scores is used to evaluate genes driving the signatures. B. Bar chart comparing the association of IIV and IMCV with Prolonged-IFN response transcriptional signatures across cell subsets. mHG was used to test enrichment on CITE-seq gene expression data. Signed -log10(adj.p) for each factor is shown. C-D. Volcano plot of genes differentially expressed between study participants with high and low factor scores in the tonic-IFN response (C) and prolonged-IFN response (D) signatures. Study participants with top and bottom 20 factor scores for IMCV are compared. Wilcoxon rank-sum test and BH correction for multiple testing were used to assess the significance of the differences. Most differentially expressed genes (adj.p <10^-4^ and |log2 fold difference| > 0.1) are annotated and colored by cell subset. E-F. Heatmap quantifying plasma cytokines relationship with IMCV of the tonic-IFN response (E) and prolonged-IFN response (F) across cell subsets. Spearman rank correlation was used to evaluate the associations of gene expression in the signatures with plasma levels of circulating cytokines. Correlations with |r|>0.25 are indicated with an asterisk. G. Boxplot showing IIV and IMCV factor scores stratified by seasonal influenza vaccine status (participants vaccinated within 6 months before sampling or not). Wilcoxon rank-sum test was performed. Significance is indicated with the following symbol: *=p<0.05.

**Figure S4.**
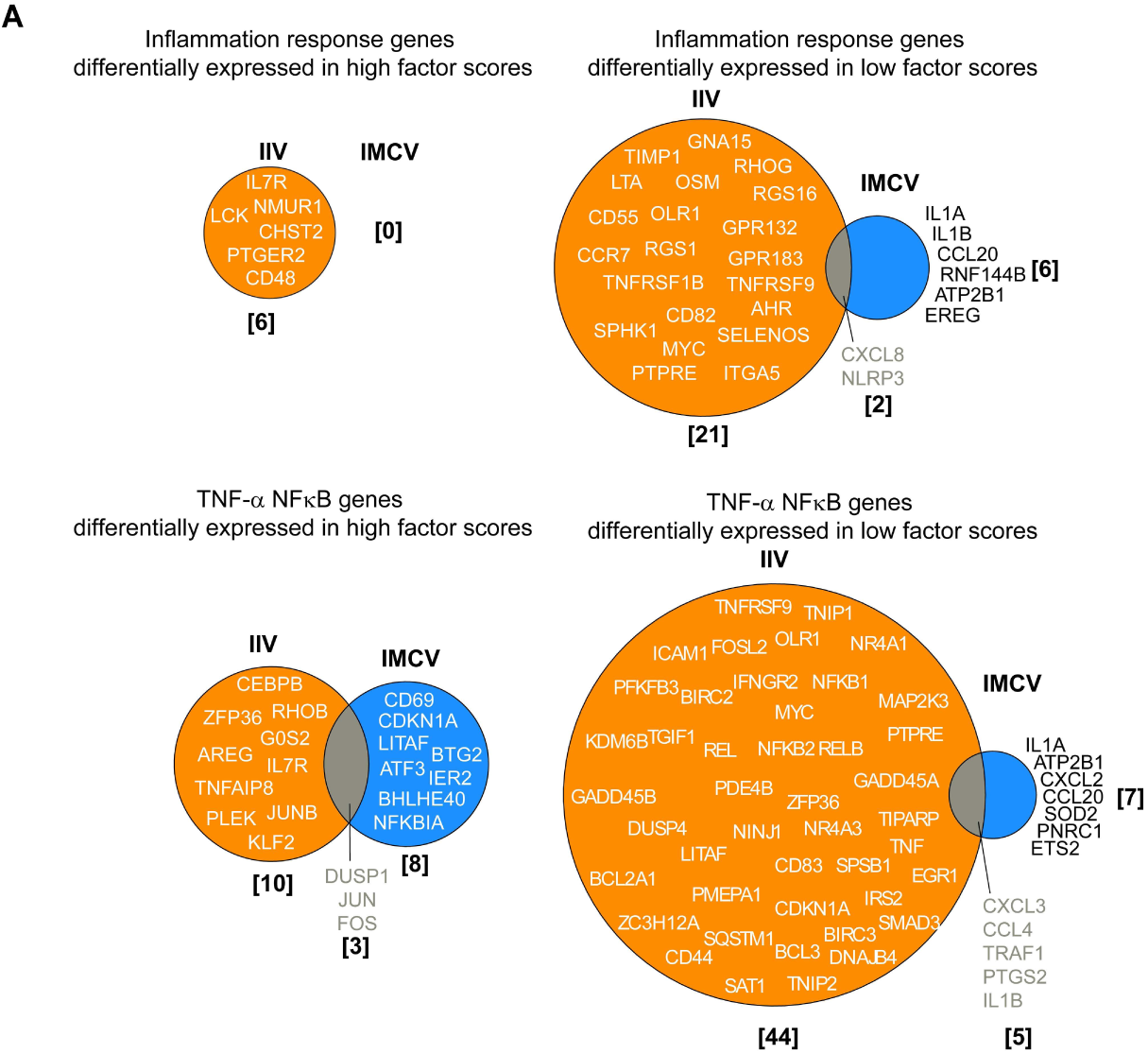
Related to Figure 4. A. Venn diagrams of differentially expressed inflammation response and TNF response genes between study participants with high (top 20%) and low (bottom 20%) IIV and IMCV factor scores. Wilcoxon rank-sum test and BH correction were used. Most differentially expressed genes (adj.p <0.1 and |log2 fold difference| > 0.25) are considered and visualized on Venn diagrams.

**Figure S5.**
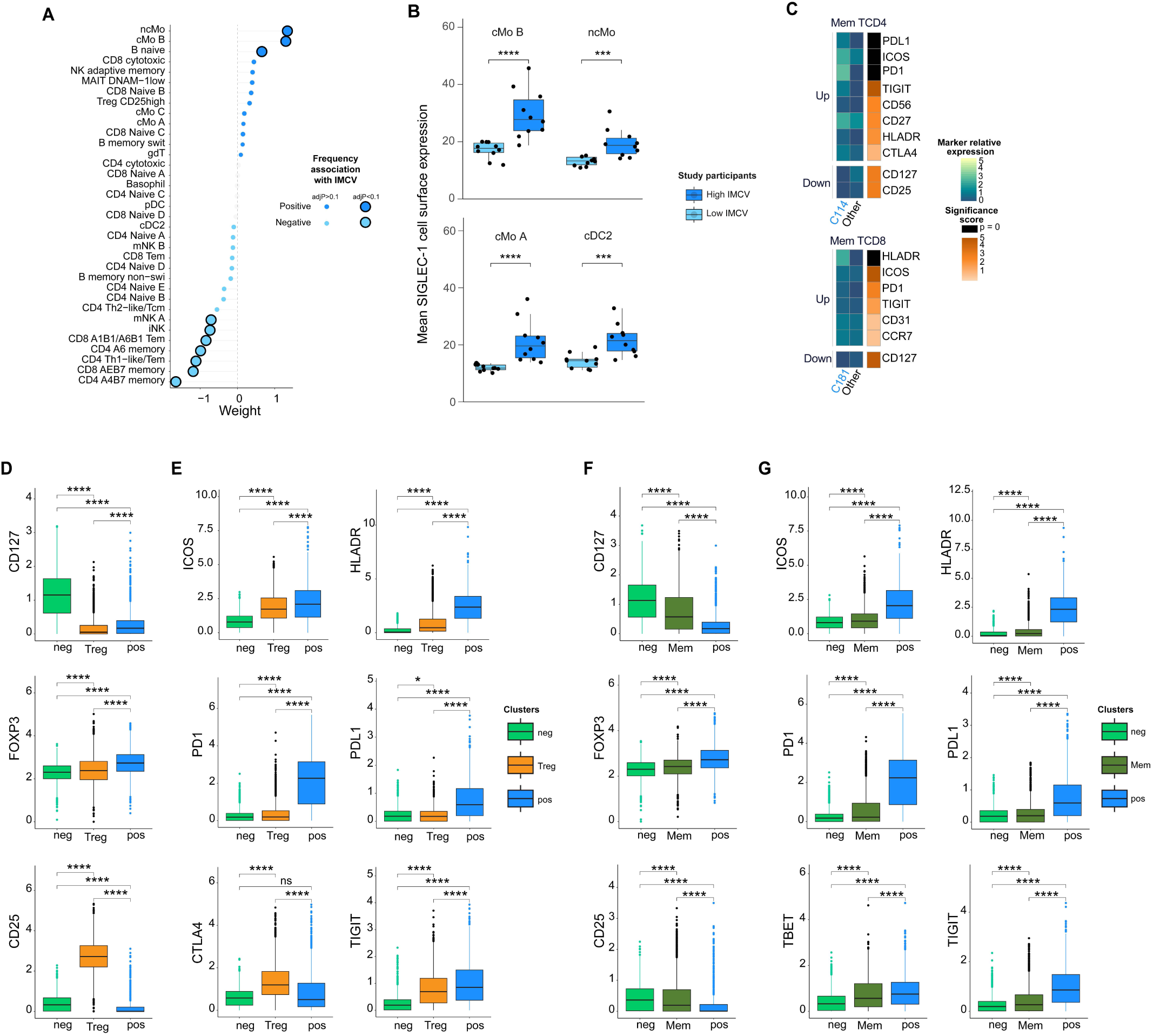
Related to Figure 5. A. Lollipop plot of CITE-Seq immune cell subset frequency association with IMCV. Large circles with black rims represent features significantly associated with IMCV (adj.p < 0.1) and small circles represent non-significant features. Linear regression was used to evaluate association of features with IMCV. Colors represent MOFA feature weight: blue indicates negative association and orange indicates positive association with IMCV. B. Box plots of SIGLEC-1 surface expression in myeloid cell subsets from individuals with high-IMCV and low-IMCV factor scores in CITE-seq. Cell subsets whose frequency is not associated with IMCV are shown. Dots indicate individual participants’ mean expression. Wilcoxon rank-sum test was used to assess significance of the difference (****=p < 0.0001, ***=p < 0.001). C. Heatmap of differential protein expression in CyTOF memory T cells. Clusters whose frequencies are positively associated with IMCV (Pos) are compared to clusters not significantly associated with IMCV (ns). Kruskal–Wallis test was used to evaluate proteins differentially expressed (p<0.00001). Post-hoc pairwise Wilcoxon rank-sum test was used to compare expression levels and medians of expression are displayed. Markers are ordered by significance using a score based on the scaled -log10(p-value), with p=0 represented in black. D-G. Boxplot showing single cell expression of memory T cell clusters whose frequencies are positively (pos) or negatively (neg) associated with IMCV, compared to those from regulatory T cells (Treg, D-E) and memory T cell clusters (Mem, F-G). Kruskal–Wallis test was used to evaluate proteins differentially expressed (p<0.00001). Canonical regulatory T cell markers (D and F) and T cell co-activation/inhibition markers (E and G) are shown. Wilcoxon rank-sum test was used to assess the significance of the differences (****=p < 0.00005, ***=p < 0.0005, **=p<0.005, *=p<0.05).

**Figure S6.**
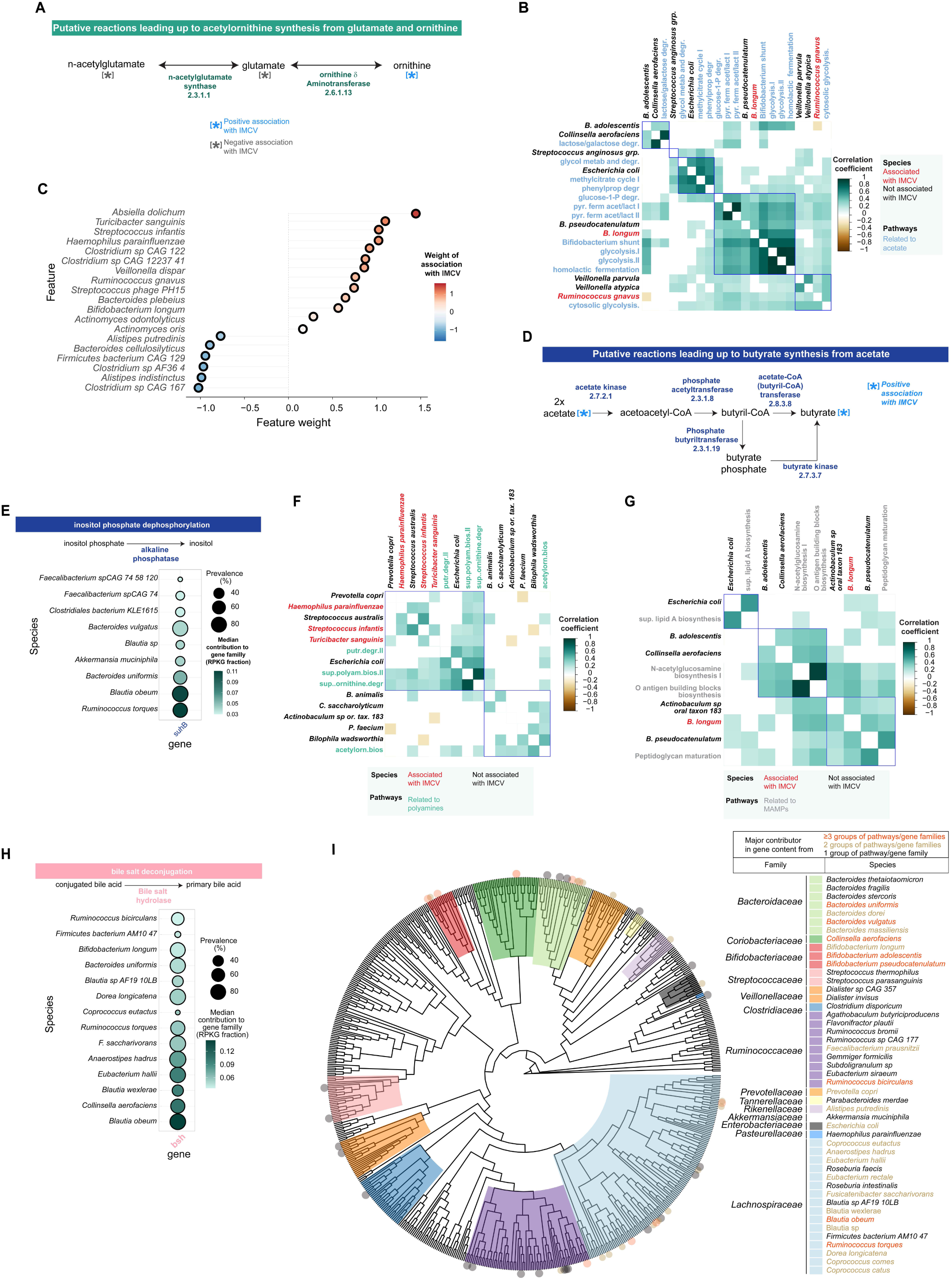
Related to Figure 6. A. Putative reactions leading up to ornithine synthesis from glutamate and n-acetylglutamate. Feature association with IMCV is indicated by an asterisk. Numbers indicate enzyme commission numbers for each reaction. B. Heatmap quantifying species relative abundances association with unstratified pathway abundances related to acetate biosynthesis. Spearman’s rank correlation was used to evaluate association (correlation coefficient > 0.4 and adj.p <0.1). Species significantly correlated with at least 1 pathway are shown. Colors indicate pathways (blue), species associated with IMCV (red) or species non-association with IMCV (black). C. Lollipop plot of microbiome species relative abundances association with IMCV. Features significantly associated with IMCV (adj.p < 0.1) are shown. Linear regression was used to evaluate association of features with IMCV. Colors represent MOFA feature weight: blue indicates negative association and orange indicates positive association with IMCV. D. Putative reactions leading up to butyrate synthesis from acetate. feature association with IMCV is indicated by an asterisk. Numbers indicate enzyme commission numbers for each reaction E. Dot plot of species contribution to the total genomic content for alkaline phosphatase gene families. Major contributors are shown (top 20 most prevalent species and median contribution > 0.025). All contributions, including from minor contributors and unclassified species contributions were documented in the Table S3. F-G. Heatmap quantifying species relative abundances association with unstratified pathway abundances related to polyamine metabolism (F) and MAMPS production (G). Spearman’s rank correlation was used to evaluate association (correlation coefficient > 0.4 and adj.p <0.1). Species significantly correlated with at least 1 pathway are shown. Colors indicate polyamine pathways (green) and MAMP pathways (gray), species associated with IMCV (red) or species non-association with IMCV (black). H. Dot plot of species contribution to the total genomic content for Bile salt hydrolase gene families. Major contributors are shown (top 20 most prevalent species and median contribution > 0.025). All contributions, including from minor contributors and unclassified species contributions were documented in the Table S3. I. Phylogenetic tree of the microbiome data colored by family. Species identified as a major contributor of gene family across the groups of metabolic processes involved in LPS endotoxin, SCFA, polyamine and primary bile acid metabolism were identified with a dot at the periphery of the tree and listed in the legend.

**Figure S7.**
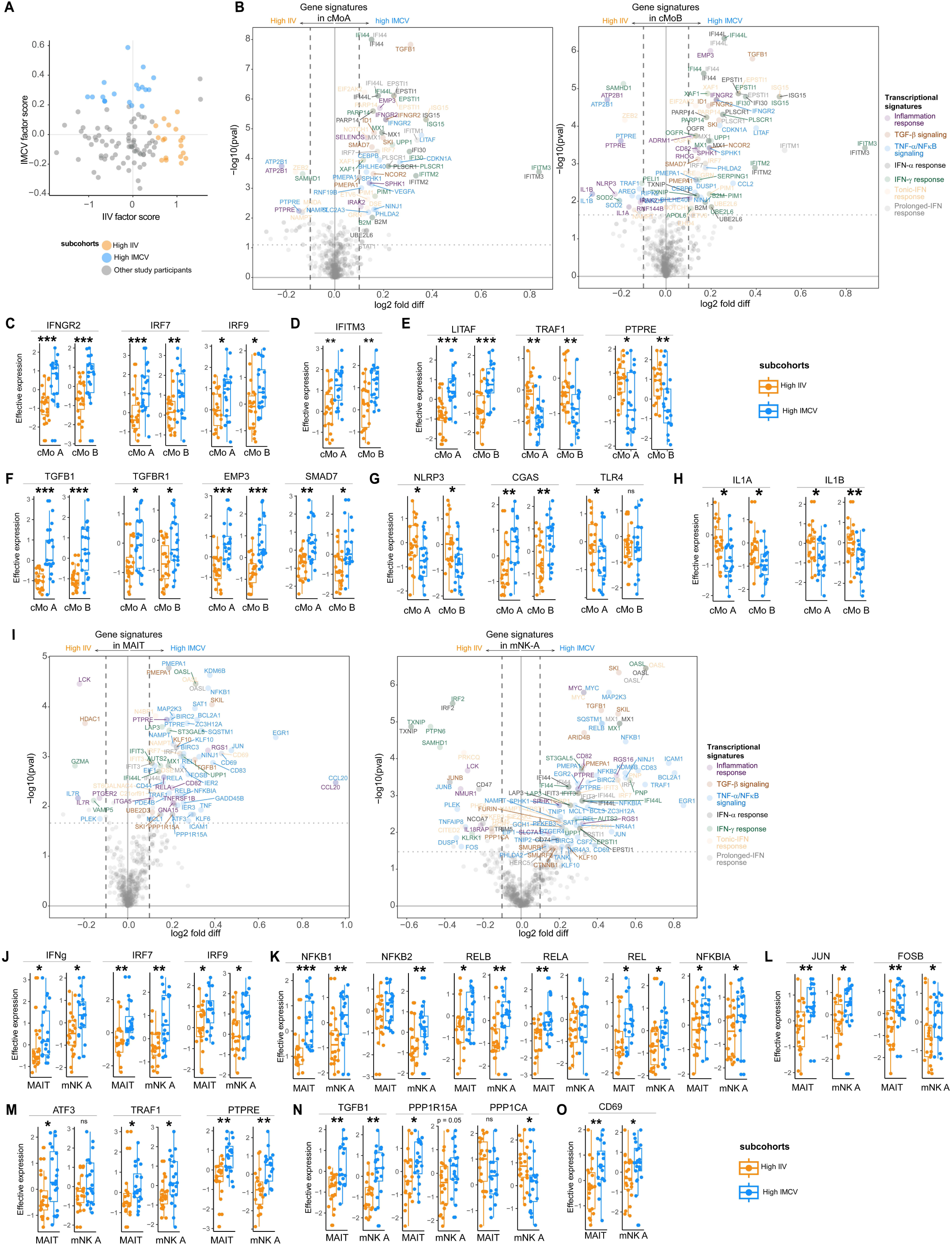
Related to Figure 7. A. Scatter plot of IIV and IMCV factor scores of ImmunoMicrobiome study participants. Colors indicate the high-IIV and high-IMCV sub-cohorts. B. Volcano plot of genes differentially expressed in cMo A and cMo B subsets between study participants of the high-IMCV and high-IIV sub-cohorts. Wilcoxon rank-sum test and BH correction for multiple testing were used to assess the significance of the differences. Most differentially expressed genes (adj.p <10^-1^ and |log2 fold difference| > 0.1) are annotated. Customized IFN-α response and IFN-γ response gene sets without overlap with inflammation signature, and vice versa, were used. C-H. Boxplot comparing expression of selective sets of genes in cMo-A and cMo-B subsets of high-IMCV and high-IIV sub-cohorts study participants including interferon pathway regulators (C), interferon stimulated genes (D), TNF-α pathway regulators (E), TGF-β pathway regulators (F), innate immune sensors (G), and IL-1 family members (H). I. Volcano plot of genes differentially expressed in MAIT and mNK A subsets between study participants of the high-IMCV and high-IIV sub-cohorts. Wilcoxon rank-sum test and BH correction for multiple testing were used to assess the significance of the differences. Most differentially expressed genes (adj.p <10^-1^ and |log2 fold difference| > 0.1) are annotated. Customized IFN-α response and IFN-γ response gene sets without overlap with inflammation signature, and vice versa, were used. J-O. Boxplot comparing expression of selective sets of genes in MAIT and mNK A subsets of high-IMCV and high-IIV sub-cohorts study participants including interferon pathway regulators (J), NFκB-related genes (K), AP-1 family members (L), TNF-α pathway regulators (M), TGF-β pathway regulators (N), and CD69 (O).

